# Safety and efficacy studies of CRISPR/Cas9 treatment of sickle cell disease in clinically relevant conditions highlights disease-specific responses

**DOI:** 10.1101/2024.01.14.575586

**Authors:** Giacomo Frati, Megane Brusson, Gilles Sartre, Bochra Mlayah, Tristan Felix, Anne Chalumeau, Panagiotis Antoniou, Giulia Hardouin, Jean-Paul Concordet, Oriana Romano, Giandomenico Turchiano, Annarita Miccio

**Affiliations:** Université Paris Cité, Imagine Institute, Laboratory of chromatin and gene regulation during development, INSERM UMR 1163, Paris, France; INSERM U1154, CNRS UMR7196, Museum National d’Histoire Naturelle, Paris, France; University of Padova, Department of Molecular Medicine, Padova, Italy; University College London, London, United Kingdom

**Keywords:** CRISPR/Cas9, genotoxicity, hemoglobinopathies, sickle cell disease, hematopoietic stem cells

## Abstract

Reactivation of fetal hemoglobin (HbF) expression through clustered regularly interspaced short palindromic repeats (CRISPR)/Cas9-mediated disruption of regulatory elements involved in γ-globin gene repression is a promising gene therapy strategy for the treatment of sickle cell disease (SCD). However, preclinical studies aimed at optimizing the genome editing process and evaluating the safety of the editing strategy are necessary to translate this approach to the clinics. This is particularly relevant in the context of SCD, a disease characterized by inflammation, which can affect hematopoietic stem and progenitor cells (HSPCs), the target cell population in gene therapy approaches for hematopoietic disorders.

Here, we describe a genome editing strategy leading to therapeutically relevant reactivation of HbF expression by targeting the binding sites (BSs) for the leukemia/lymphoma related factor (LRF) transcriptional repressor in the *HBG1* and *HBG2* γ-globin promoters. Electroporation of Cas9 ribonucleoprotein and single guide RNA (sgRNA) targeting the *HBG* promoters in healthy donor (HD) and patient-derived HSPCs resulted in a high frequency of LRF BS disruption and potent HbF synthesis in their erythroid progeny differentiated *in vitro* and *ex vivo* after transplantation into immunodeficient mice. LRF BS disruption did not impair SCD and HD HSPC engraftment and differentiation, but was more efficient in SCD than in HD cells. However, SCD HSPCs showed a reduced engraftment and a myeloid bias compared to HD cells.

Importantly, in HSPCs, we detected off-target activity and the intra- and inter-chromosomal rearrangements between on- and off-target sites, which were more pronounced in SCD samples (likely because of the higher overall editing efficiency), but did not impact the target gene expression. Off-target activity was observed *in vitro* and *in vivo*, thus indicating that it does not impair engraftment and differentiation of SCD and HD HSPCs. Finally, transcriptomic analyses showed that the genome editing procedure results in the upregulation of genes involved in DNA damage and inflammatory responses in both HD and SCD samples, although gene dysregulation was more evident in SCD HSPCs.

Overall, this study provides evidences of feasibility, efficacy and safety for a genome editing strategy based on HbF reactivation and highlights the need of performing safety studies, when possible, in clinically relevant conditions, i.e., in patient-derived HSPCs.

## Introduction

Sickle cell disease (SCD) is an autosomal recessive inherited blood disorder caused by a single mutation in the β-globin gene (*HBB*) leading to an amino acid substitution (β7^Glu>Val^) in the β-globin chain and resulting in the production of an abnormal hemoglobin (hemoglobin S; HbS), which polymerizes under deoxygenated conditions. This ultimately leads to the formation of distorted “sickle shaped” red blood cells (RBCs) that can occlude small vessels. The clinical phenotype of patients affected by SCD is mainly characterized by vaso-occlusive crises (VOCs), hemolysis, iron overload and multi-organ damage. It has been estimated that between 300,000 and 400,000 neonates affected by SCD are born each year, the majority of these births occurring in sub-Saharan Africa^1^. Patients affected by SCD are usually treated with palliative treatments consisting of lifelong RBC exchange transfusions and iron chelation, in order to decrease the frequency of VOCs, suppress anemia and reduce iron toxicity. These therapies significantly improve their quality of life, but do not provide a definitive cure for SCD patients. Allogeneic hematopoietic stem cell transplantation (HSCT) is a curative option, but it is limited by the availability of compatible human leukocyte antigen (HLA)-matched donor and the potential graft rejection and graft-versus-host diseases^2^.

Decades of advances in gene therapy resulted in the development of several strategies based on the transplantation of genetically modified autologous HSCs, thus overcoming the limitations associated with allogeneic HSCT. Initially, gene therapy strategies were based on the transplantation of lentiviral (LV)-engineered HSCs harboring a functional^3,4^ and anti-sickling^5,6^ *HBB* transgene. The overall outcomes of these studies are encouraging since most of the recruited patients became transfusion-independent. Nevertheless, a number of issues, such as the lack of an optimal protocol for HSC transduction, the low β-globin expression per vector copy and the poor engraftment of transduced HSCs, still remain unsolved^7^. Furthermore, the use of LVs is intrinsically associated with the risk of genotoxicity due to their semi-random integration (mainly in the genes’ body). Recently, the occurrence of malignant transformations has been reported in two SCD patients who underwent an LV-based clinical trial^8,9^, causing a partial clinical hold in Europe and the US, although no causal link between these events and LV transduction has been proven.

The introduction of genome editing (GE) technologies based on designer nucleases allowed the development of novel and safer strategies for the treatment of SCD. Due to its high efficacy and accessibility, the clustered regularly interspaced short palindromic repeats (CRISPR)/Cas9 nuclease system has emerged as a prominent tool to manipulate the genome. Nuclease-based GE strategies for the treatment of SCD include (i) the correction of disease-causing mutation and (ii) the reactivation of γ-globin genes (*HBG1/2*) that are normally silenced soon after birth^10^, as a genetic condition causing hereditary persistence of fetal hemoglobin (HPFH) in adulthood ameliorates the SCD clinical phenotype^11^. If correction of the SCD-causing mutation held the promise to precisely and definitively cure the disease, its clinical translation might be hampered by the low efficacy of homology directed repair (HDR) pathway in HSCs^12–16^, the DNA repair mechanism required in gene correction approaches. On the contrary, γ-globin reactivation can be achieved by the CRISPR/Cas9-mediated manipulation of several cis-regulatory elements, which are easily and efficiently inactivated by exploiting the non-homologous end joining (NHEJ) pathway that is highly active in HSCs. The target cis-regulatory elements include genomic regions modulating the expression of γ-globin transcriptional repressors^17–19^ and their binding sites within the *HBG1/2* promoters^20–22^. By targeting the *HBG1/2* promoters, we have shown that it is possible to evict the leukemia/lymphoma related factor (LRF; also known as ZBTB7A or FBI-1), a known repressor of *HBG* expression and reactivate HbF^22^.

Regardless of the specific strategy, a number of concerns related to the safety profile of designer nucleases recently emerged. In fact, nuclease-mediated DNA double-strand breaks (DSBs) formation is the source of several unwanted events^23^. Cas9-sgRNA treatment of human HSPCs induces a DNA damage response that can lead to apoptosis^24,25^. CRISPR/Cas9 can cause P53-dependent cell toxicity and cell cycle arrest, resulting in the negative selection of cells with a functional P53 pathway^26^. Furthermore, the generation of DSBs at on- and/or off-target sites is associated with the risk of large genomic rearrangements, such as deletions, inversions and translocations^27–31^. The potential toxicity of the GE procedure might be exacerbated in SCD HSPCs. In fact, although clinically heterogeneous, SCD is a chronic inflammatory disease^32^. The incidence of nearly every clinical manifestation of SCD correlates with high white blood cell count, indicating a role for leukocytes and inflammation in the pathophysiology of SCD. Leukocytosis is common in SCD patients and is manifested by elevation in monocyte and neutrophil counts^33–35^, accompanied by elevated levels of circulating inflammatory cytokines, including tumor necrosis factor a (TNF-a), interleukin (IL)-1, and IL-8^36^. The difficulty in obtaining high cell doses and a robust engraftment of LV-transduced HSPCs in SCD patients suggests that the unique inflammatory bone marrow (BM) environment associated with SCD may have a significant impact on HSPCs^37^. Furthermore, a recent study in a mouse model of SCD showed that HSPCs are characterized by a high mutational burden and a high incidence of clonal hematopoiesis (potentially enhanced by the inflammation environment) and leukemias have been reported in SCD patients^38–40^. For these reasons, the safety profile of gene therapy approaches must be carefully evaluated in patients’ HSPCs.

In this study, we analyzed the effects of CRISPR/Cas9-mediated GE in a side-by-side comparison between HD- and SCD patient-derived HSPCs to evaluate the efficacy and safety of a gene therapy approach targeting the *HBG1/2* promoters in clinically relevant conditions.

## Results

### Optimization of LRF BS editing via Cas9 RNP delivery in HSPCs induces HbF expression in their erythroid progeny

We previously edited human SCD HSPCs by electroporating RNP complexes containing Cas9 and sgRNAs (−196 or −197 sgRNA) targeting the LRF BS in the −200 region of the *HBG1/2* promoters; LRF eviction led to a potent γ-globin reactivation^22^ (**Figure 1A**). However, the GE procedure was associated with a considerable toxicity in adult HSPCs^22^.

**Figure 1.**
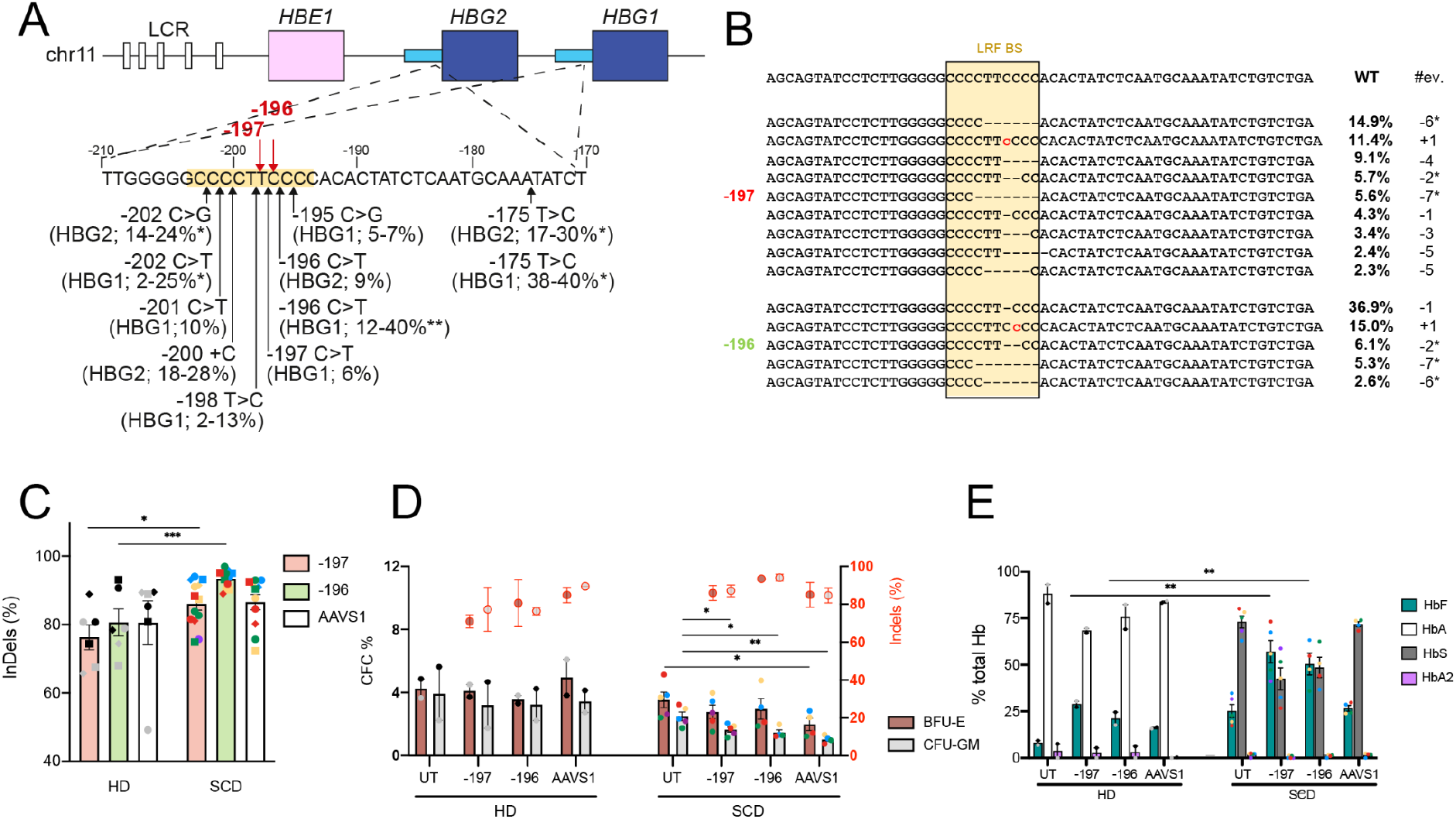
Targeting the LRF binding site in the *HBG* promoters is highly efficient and reactivates HbF expression. (**A**) Schematic representation of the β-globin locus on chromosome 11, depicting the hypersensitive sites (white boxes) of the locus control region (LCR) and the *HBE1*, *HBG2*, and *HBG1* genes (colored boxes). The sequence of the *HBG2* and *HBG1* identical promoters (from −210 to −170 nucleotides upstream of the *HBG1/2* TSS) is shown below. Black arrows indicate HPFH mutations described at *HBG1* and/or *HBG2* promoters, with the percentage of HbF in heterozygous carriers of HPFH mutations^22^. The highest HbF levels were generally observed in individuals carrying SCD (*) or β-thalassemia mutations (**). The LRF BS is highlighted by a yellow box. Red arrows indicate the −196 and −197 sgRNA cleavage sites. (**B**) NGS sequencing of the *HBG1/2* promoters treated with the −197 and −196 sgRNAs as compared with the wild-type sequence (WT). The analysis shows the percentage of individual InDel events identified in mature erythroblasts derived from adult SCD HSPCs (n=4)^22^. #ev. and * indicate the type of InDel (i.e., - for deletions, + for insertion) and the deletions associated with microhomology motifs, respectively. (**C**) Bar plot showing InDel frequency in cells treated with the −197, −196 and AAVS1 sgRNAs [e.g., in undifferentiated HSPCs (circles), burst forming units-erythroid colonies (BFU-E, squares) and granulocyte–macrophage progenitor (CFU-GM, rhombuses) of healthy donor (HD) and sickle cell disease (SCD) cells]. *p<0.05 and **p<0.01; unpaired t-test. (n= 2 donors for HD; n=5 donors for SCD). (**D**) Frequency of CFCs in healthy donor (HD) and sickle cell disease (SCD) HSPCs transfected with Cas9 RNP. Untreated samples (UT) served as control. Red-circled dots indicate InDels values at the *HBG* promoters specifically measured in CFC populations (see right Y-axis). Data are expressed as mean ± SEM (n= 2 donors for HD; n=5 donors for SCD). Donors are indicated using different colors. Samples derived from the mobilized SCD patients are indicated in red. *p<0.05; **p<0.01 Unpaired t-test. (**E**) Hemoglobin expression in HD and SCD BFU-E, as measured by CE-HPLC. **p<0.01; unpaired t-test. (n= 2- 5 donors). We used 4 non-mobilized and one plerixafor-mobilized SCD HSPC samples; 2 G-CSF-mobilized HD HSPC samples. In **C** and **D** each color represents a different donor. Samples derived from the mobilized SCD patients are indicated in red.

First, to optimize HSPC fitness while maintaining a high GE efficiency, we tested different parameters such as different Cas9 variants, a transfection enhancer and small molecules known to preserve stemness. The use of a Cas9 with two SV40 nuclear localization signals (NLSs; one at the N- and one at the C-terminus of Cas9) fused or not to GFP led to similar editing efficiencies in cord-blood (CB)-derived HD CD34^+^ HSPCs using an sgRNA targeting the *HBG* promoters (−197 sgRNA^22^; **Supplementary Figure 1A**). The addition of a third NLS (the myc NLS at the C-terminus of Cas9) did not improve GE efficiency (**Supplementary Figure 1A**). The addition of the transfection enhancer substantially increased GE efficiency (**Supplementary Figure 1A**). Editing efficiency increased over time, indicating persistent Cas9 cleavage activity for several days after transfection (**Supplementary Figure 1A**). However, the up-regulation of the *CDKN1A* gene (a downstream effector of P53) peaked at 15-24 hours after transfection, but was transient and *CDKN1A* expression returned to normal levels after two to three days (**Supplementary Figure 1B**). The Cas9x2NLS_GFP variant was used in the following experiments because of its good efficiency and the presence of the GFP that allows the monitoring of transfection efficiency. We also used the transfection enhancer, which substantially increased GE efficiency without increasing cell toxicity.

We then edited and cultured CB-derived HD and adult peripheral blood, non-mobilized SCD HSPCs in presence or in absence of two small molecules (i.e. SR1 and UM171) known to preserve the HSC stemness^41^. Insertion and deletion (InDel) frequency was unchanged upon treatment with each of these compounds (although the combination tended to increase GE efficiency in HD samples), and was significantly higher in SCD than in HD samples (**Supplementary Figures 1C**). The addition of SR1 and UM171 did not reduce cell mortality typically observed after transfection, particularly in adult SCD samples (**Supplementary Figure 1D**). In parallel, we performed a time course analysis of different cell populations with increasing stemness properties. Importantly, control and CRISPR/Cas9 electroporated cells showed a comparable number of the different subpopulations over time, confirming that the editing procedure does not have an impact (e.g., cell death) on specific cell subpopulations (**Supplementary Figure 1E**). Interestingly, at day 6 the number of CRISPR/Cas9 electroporated cells in the medium without UM171 and SR1 was substantially reduced compared to non-edited cells, suggesting a detrimental effect of the treatment on cell proliferation. However, the combined use of UM171 and SR1 rescued the cell number defect observed upon GE (**Supplementary Figure 1E**).

We then applied this protocol in a side-by-side comparison between adult SCD and HD HSPCs and evaluated the effects of LRF BS editing on γ-globin expression. HSPCs were treated with Cas9 RNP and the −197 or the −196 sgRNAs targeting the LRF BS in the *HBG1/2* promoters (**Figure 1A**). A sgRNA targeting the unrelated *AAVS1* region was used as control. As previously reported^22^, *HBG*-targeting sgRNAs efficiently destroy the LRF BS by producing different InDel profiles. In particular, the sgRNA −196 mainly produces 1-bp InDels while the sgRNA −197 produces 4- to 7-bp deletions. Both *HBG*-targeting sgRNAs produce, at different frequencies, deletions associated to microhomology (MH) motifs (i.e. −7*, −6*, −2*). These events could be generated by microhomology-mediated end joining (MMEJ), which takes place through annealing of short stretches of identical sequence flanking the DSB but is thought to be less active in long-term repopulating HSCs compared to NHEJ^19,42^ (**Figure 1B and Supplementary Figure 2A**). As previously observed (**Supplementary Figure 1D)**, we detected a significant higher InDel frequency in SCD compared to HD samples (**Figure 1C**). However, the InDel profile was similar between HD and SCD cells (**Supplementary Figure 2A**). A colony forming cell (CFC) assay showed a decreased clonogenic potential for SCD compared to HD HSPCs upon CRISPR/Cas9-mediated editing (**Figure 1D**).

LRF BS editing resulted in a robust HbF expression as measured by CE-HPLC in erythroid colonies (**Figure 1E**). Interestingly, SCD samples showed a higher basal HbF expression, and upon *HBG* editing, HbF levels exceeded HbS and were overall higher compared to HD samples, suggesting that *HBG* promoter priming allows a more efficient HbF reactivation in patients’ cells (**Figure 1E**). Interestingly, we observed a significantly higher ^A^γ (*HBG1*) production compared to ^G^γ (*HBG2*; **Supplementary Figure 2B**). This is likely due to the simultaneous targeting of *HBG1* and *HBG2* promoters causing the loss of the 4.9-kb region containing the *HBG2* gene (**Supplementary Figure 2C**).

### LRF BS editing is maintained in long-term repopulating HSCs

To demonstrate the editing of repopulating HSCs, we transplanted untreated (UT) and Cas9-treated CD34^+^ HSPCs into immunodeficient mice (4 non-mobilized and one plerixafor-mobilized SCD HSPC samples; 2 G-CSF-mobilized HD HSPC samples). 16 weeks after transplantation, the chimerism was significantly higher in mice transplanted with HD cells (36.16 ± 4.72% and 25.35 ± 4.26% in BM and spleen, respectively) than in animals transplanted with SCD cells (5.48 □ 1.85% and 1.34 □ 0.14% in BM and spleen, respectively) including those transplanted with plerixafor-mobilized SCD HSPCs (**Figure 2A**). However, CRISPR/Cas9 editing had no impact on HD HSC engraftment, while a mild reduction was observed in 3 out of 5 SCD donors (**Figure 2A**). Importantly, LRF BS editing did not affect multi-lineage differentiation in neither HD nor SCD samples (**Figure 2B and Supplementary Figure 3A**). Interestingly, compared to HD cells, patient-derived cells showed a higher propensity to differentiate toward the CD14^+^ or CD11b^+^ myeloid lineage in the BM (**Figure 2B**).

**Figure 2.**
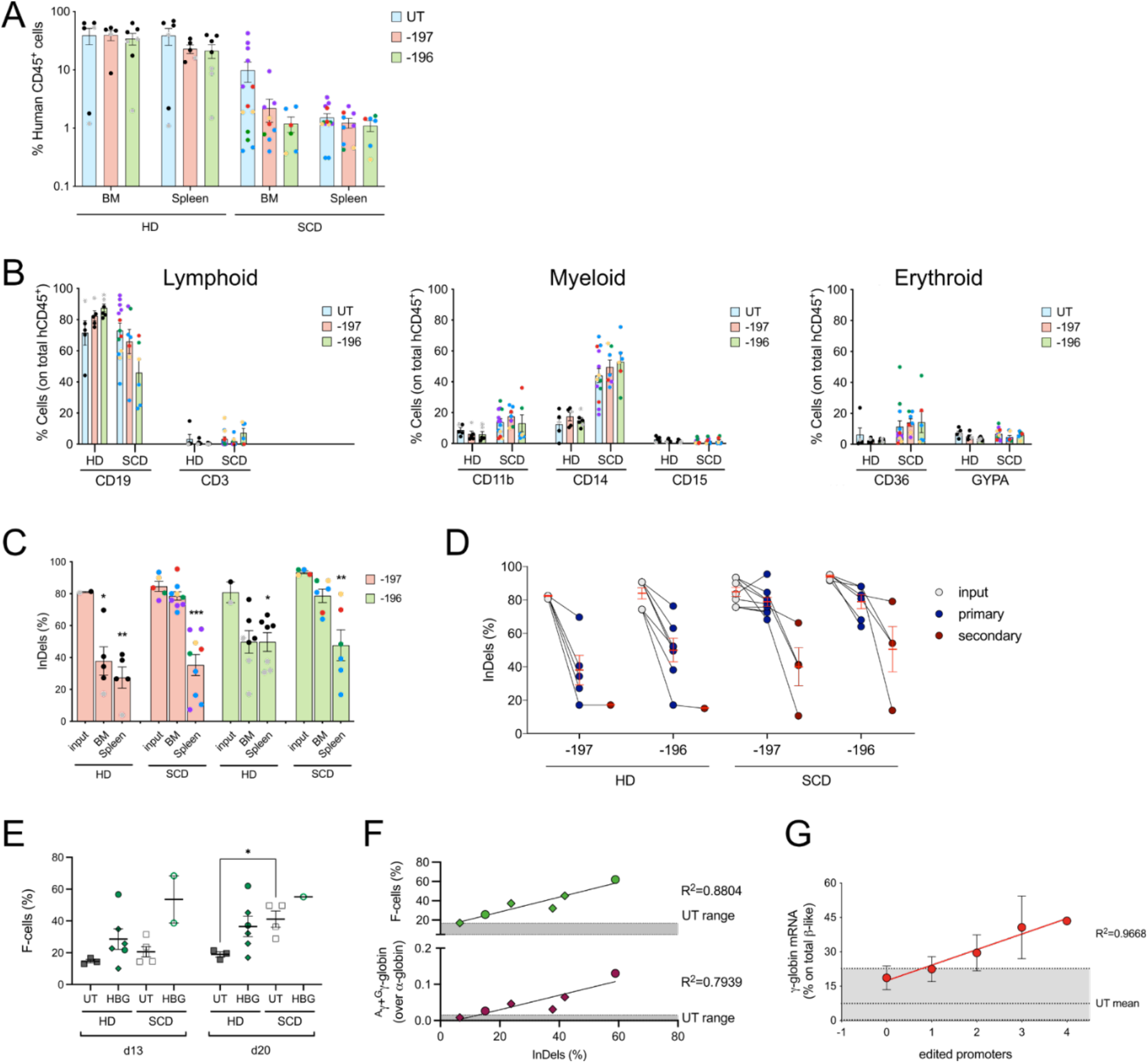
Editing of bona fide HSCs leads to HbF reactivation in their erythroid progeny. (**A**) Engraftment in NSG mice transplanted with untreated (UT) and edited HD or SCD human HSPCs 16 weeks after transplantation. Engraftment is represented as the percentage of human CD45^+^ cells in the total murine and human CD45^+^ cell population in bone marrow (BM) and spleen. Values shown are means ± SEM. (**B**) Frequency of human T (CD3) and B (CD19) lymphoid, myeloid (CD14, CD15 and CD11b) and erythroid (CD36, GYPA) in the BM of mice transplanted with control and edited HSPCs. (**C**) Editing efficiency in the input cells and in BM- and spleen-derived human CD45^+^ progeny of repopulating HSCs. GE frequency was evaluated by Sanger sequencing and TIDE analysis. Values shown are means ± SEM. Asterisks indicate statistical significance between input and repopulating cells in each group (*p<0.05; **p<0.01; ***p<0.001; unpaired t-test). Each data point represents an individual mouse; each color identifies a different donor. (**D**) InDel frequency in secondary recipient mice (red dots) as compared to the corresponding primary recipient (blue dots) and input cells (grey dots). Red bars indicate means ± SEM. (**E**) Percentages of HbF- expressing cells (F-cells) in UT and *HBG*-edited [with the −197 (circles) or −196 (rhombuses) sgRNAs] samples derived from HD (filled dots) and SCD (empty dots) cells. (**F**) XY graph showing the correlation between the % of F-cells (upper panel) or the amount of ^A^ γ- and ^G^ γ-globin chains (as determined by RP-HPLC, lower panel) and the InDel frequency at the *HBG* promoters in HD erythrocytes edited with the sgRNA −197 (circles) and −196 (rhombuses). (**G**) Correlation between the γ-globin mRNA expression and the number of edited *HBG* promoters in BFU-E from repopulating HD HSPCs treated with the −197 sgRNA (n=31 individual colonies).

The InDel frequency at the *HBG1/2* promoters in HD cells treated with the −197 or −196 sgRNAs was decreased in human CD45+ cells repopulating the BM and the spleen compared to the input cells (**Figure 2C**). Interestingly, we measured a higher InDel frequency in human cells engrafting the BM of mice transplanted with SCD cells compared to those who received HD HSPCs (**Figure 2C-D**). Importantly, the InDel spectrum in *HBG*-edited cells was largely similar before and after transplantation in mice receiving HD and SCD HSPCs with no evidence of clonal dominance *in vivo* (**Supplementary Figure 3B**). All the MH-associated events except the 6-bp deletion (−6*) in samples treated with the −196 sgRNA were detected in human cells repopulating both primary and secondary recipient animals. These results indicate that the −6* event occurs *in vitro* mostly in progenitors through MMEJ, a pathway that is disfavored in long-term repopulating HSCs *in vivo*. However, this event represents a minimal proportion of the total InDels *in vitro* and cannot account for the reduction in GE efficiency *in vivo* in mice receiving HD cells (**Supplementary Figure 3B**).

### *HBG* promoter editing induces HbF expression in the erythroid progeny of BM repopulating cells

Since the NSG mouse model does not support a complete human erythroid differentiation, we evaluated the efficacy of our strategy in the erythroid populations *ex vivo* differentiated from BM repopulating human CD45^+^ cells obtained from mice showing a high chimerism (mainly animals receiving HD HSPCs). The immunophenotypic time course analysis of erythroid specific markers in erythroid liquid cultures revealed no obvious differences between UT and *HBG*-edited cells (**Supplementary Figures 4A-B**). Furthermore, we obtained a large fraction of mature enucleated RBCs from both edited HD and SCD cells (**Supplementary Figure 4A**), thus indicating that the CRISPR/Cas9 treatment does not impair erythroid differentiation. LRF BS editing resulted in an increased percentage of HbF expressing cells (F-cells) (**Figure 2E**). The proportion of F-cells and the amount of γ-globin chains measured in HD erythrocytes was positively correlated with the InDel frequency (**Figure 2F**). Similarly, in erythroid colonies (BFU-E) obtained from BM human CD45^+^ cells edited using the −197 gRNA, γ-globin mRNA levels positively correlated with the number of edited *HBG* promoters (**Figure 2G**).

### CRISPR/Cas9 leads to unwanted off-target activity and chromosomal rearrangements

We then evaluated the occurrence of unwanted genetic changes in primary human hematopoietic cells from HDs and SCDs *in vitro* (i.e. input cells) and, when possible, *in vivo* (human CD45^+^ cells isolated from the BM of NSG mice). We observed >50% of 4.9-kb deletions caused by the simultaneous cleavage of the two *HBG* promoters in both HD and SCD input cells (**Figure 3A**). Of note, these deletions were detected also in human CD45^+^ cells repopulating immunodeficient mice, suggesting that they do not impair the engraftment and multilineage differentiation of HSPCs. As observed for the InDel frequency (**Figure 2B**), the percentage of 4.9-kb deletions was maintained in SCD repopulating cells *in vivo*, while it decreased in HD samples (**Figure 3A**).

**Figure 3.**
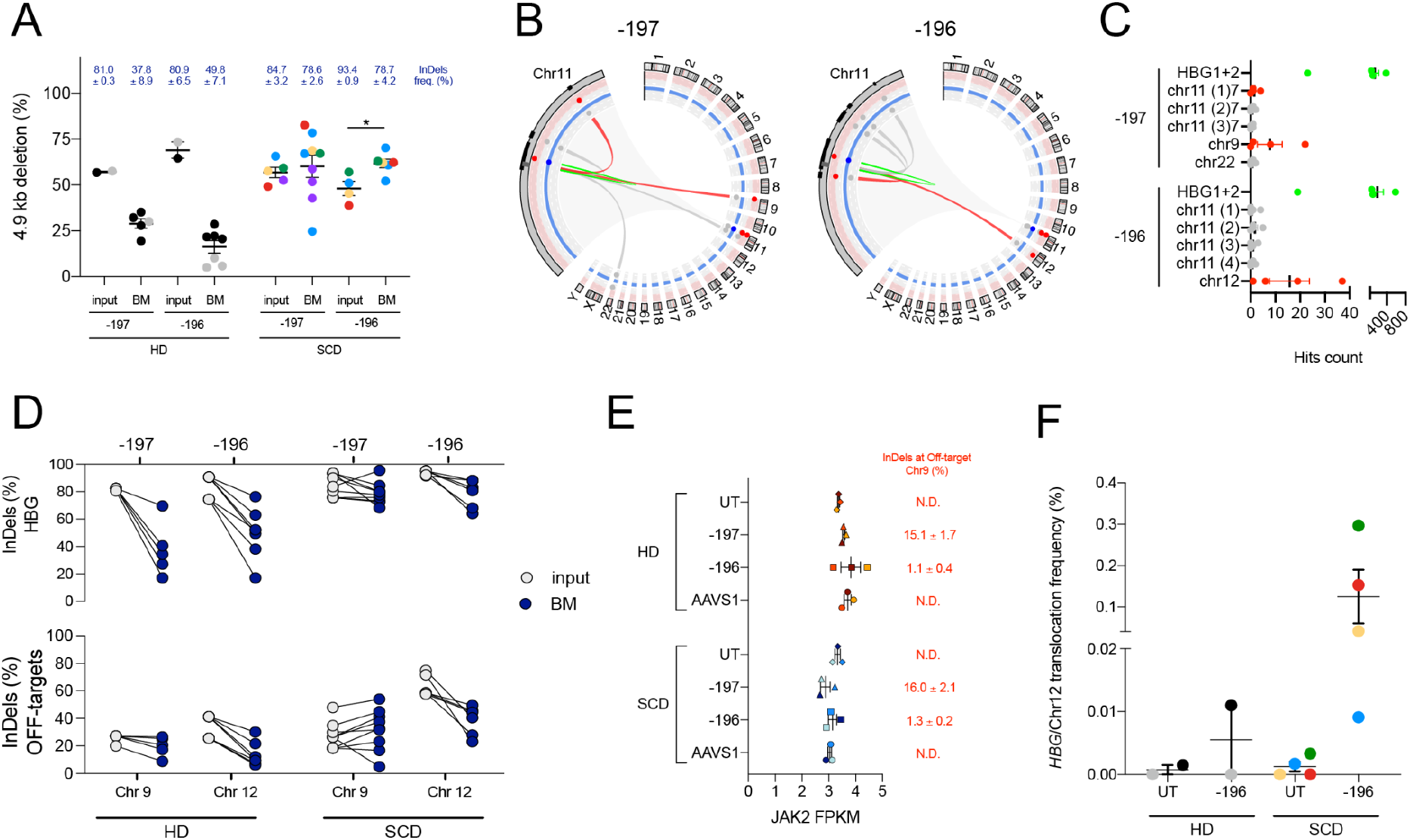
Analysis of off-target activity and large genomic rearrangements in primary HSPCs. (**A**) Frequency of 4.9-kb deletion measured by ddPCR in DNA samples from input cells and BM-repopulating human CD45^+^ cells derived from HD and SCD HSPCs. Each color identifies a donor (**p<0.01; ****p<0.0001, unpaired t-test). (**B**) Visualization of chromosomal rearrangements in −197- and −196-treated HD HSPCs by CAST-seq analysis. The Circos plot shows mutations stemming from the two on-target sites on chromosome 11 (zoomed in the left side portion). From the outer to the inner layer, black rectangles show the DNA cluster location of mutations close the on-target site; pail red and blue circles represent the threshold levels utilized to determine, respectively, the gRNA homology and the nearby sequence homology calculated in this analysis to categorize the mutation cause. The connectors represent mutations occurred by on-target cleavage activity (green), off-target cleavage activity (red), homology mediated mutations (blue) and natural break sites (grey) when no homologies with gRNA or nearby sequences are found. (**C**) Summary of putative aberrations with their relative quantification (expressed as number of hit count– run in triplicate), as determined by CAST-seq in HSPC treated with the −197 and −196 sgRNAs. Green dots indicate pooled (*HBG1* + *HBG2*) on-targets related mutations. Grey dots indicate mutations caused by naturally occurring DSBs. Red dots indicate mutations caused by off-targets. (**D**) Comparison between the InDel frequency at the on-target sites (*HBG*, upper panel) and off- target sites (indicated below, bottom panel) in transplanted HSPCs (input) and BM-repopulating human CD45^+^ cells (BM) from HD and SCD cells treated with the −197 or −196 sgRNAs (***p<0.001, unpaired t-test). (**E**) JAK2 expression as determined by RNA-seq in plerixafor-mobilized HD or SCD HSPCs treated with the −197, the −196 or the AAVS1 sgRNA. (**F**) Frequency of *HBG*/Chr12 translocation as measured by ddPCR in DNA samples from *in vitro* cultured HD and SCD HSPCs.

In our previous work, we identified putative off-target sites generated by the −196 and −197 sgRNAs using GUIDE-seq in the human kidney embryonic HEK293 cell line. This method, however, is poorly efficient in primary human HSPCs and off-targets nominated using GUIDE-seq in HEK293T cell are often not validated in HSPCs, likely because of the different chromatin structure between primary hematopoietic cells and HEK293. Therefore, we exploited CAST-seq^30^, a newly established high throughput technique for evaluating off-target activity and chromosomal rearrangements in human primary HSPCs.

CAST-seq identified several putative off-targets of the −196 and −197 sgRNAs in HD CD34^+^ cells (**Figure 3B and 3C; Supplementary Tables 1 and 2**). Amongst them, we identified one off-target mapping to the third intron of the *JAK2* gene on chromosome 9 and one off-target mapping to an intergenic region on chromosome 12 in cells treated with the sgRNAs −197 and −196, respectively (**Figure 3B-C**). These findings were confirmed by measuring the InDel frequency at the two off-target sites in both input populations and engrafted human cells (**Figure 3D**). Off-target activity was observed both *in vitro* and *in vivo* with a frequency that parallels that observed at the on-target site, suggesting that these events do not impair the engraftment and differentiation capability of HSPCs (**Figure 3D**). Importantly, *JAK2* expression was similar in control and treated samples (**Figure 3E**).

Furthermore, CAST-seq showed the occurrence of a chromosomal translocation between the *HBG1/2* promoters and off-target sites of each *HBG*-targeting sgRNA (**Figure 3B-C; Supplementary Tables 1 and 2**). A PCR performed using primers surrounding the *HBG*/Chr12 translocation junction resulted in the amplification of a specific band for the sample treated with the sgRNA −196 (**Supplementary Figure 5A**). Sanger sequencing confirmed the occurrence of the translocation induced by the sgRNA −196 (**Supplementary Figure 5B**). On the contrary, PCR designed to detect *HBG*/Chr9 translocation in samples treated with sgRNA −197 did not result in amplification of any specific product. By ddPCR, we measured a frequency of *HBG*/Chr12 translocation of 0.0055 ± 0.0055 % and 0.1250 ± 0.0079 % in −196 sgRNA-treated HSPCs from HD and SCD, respectively (**Figure 3F**). Of note, in both −196 and −197 sgRNA-treated HSPCs, we also observed other large genomic rearrangements including translocations between the *HBG1/2* promoters and likely natural occurring DSBs or potential deletions within chromosome 11 (**Supplementary Tables 1 and 2).**

### CRISPR/Cas9-mediated editing induces more prominent transcriptomic changes in SCD cells

To investigate the transcriptomic changes associated with the CRISPR/Cas9-mediated targeting of the *HBG* promoters in HD and SCD cells, and assess the safety profile of this strategy, we analyzed the global gene expression profile of HSPCs obtained from age- and sex-matched HDs and SCD patients. Both SCD patients and HDs underwent plerixafor-mediated mobilization regimens before cell harvesting, as currently performed in gene therapy protocols for SCD patients. HSPCs were preactivated for two days and then electroporated with Cas9 RNP. Two days after the CRISPR/Cas9 treatment, the total RNA was extracted and analyzed to evaluate the transcriptome of the potential drug product (genetically modified HSPCs typically injected two days after the treatment).

Comparing the transcriptome of the SCD *vs* the HD untreated (UT) samples, we observed few differentially expressed genes (DEGs; FDR<0.05 and FC>|2|), of which 41 and 60 were up- and down-regulated, respectively (**Supplementary Table 3**). These data indicate that, after four days of culture in the presence of HSPC-supporting cytokines, untreated SCD and HD cells show a similar transcriptome.

Then we evaluated the impact of CRISPR/Cas9-mediated treatment via electroporation on the HSPC gene expression profile. SCD and HD cells were treated with −197 and −196 *HBG*-targeting sgRNAs and the control AAVS1 sgRNA. Despite some variability in the HD samples, SCD cells tend to show higher levels of genome editing at both *HBG1/2* and *AAVS1* loci (**Figure 4A**). Principal component analysis (PCA) revealed a distinct expression profile associated to the donor (PC2) and to the CRISPR/Cas9 treatment (PC1) but independently from the specific sgRNA (**Figure 4B**). We observed 360, 336 and 470 upregulated genes and 45, 28 and 52 downregulated genes in HD samples treated with −197, −196 and AAVS1 sgRNA, respectively, compared to UT cells (FDR<0.05 and FC>|2|; **Supplementary Table 4**). Amongst them, 281 up- and 22 down-regulated genes were shared in all CRISPR/Cas9-treated samples regardless of the sgRNA used (**Supplementary Figure 6A**). This suggests that a high fraction of the DEGs was dysregulated by the transfection procedure and DSB formation rather than by the targeting of a specific locus. Interestingly, the number of up- and down-regulated genes in CRISPR/Cas9-treated vs UT SCD HSPCs was higher compared to that observed in HD samples. In particular, we observed 508, 599 and 737 upregulated genes and 69, 88 and 94 downregulated genes in SCD samples treated with the sgRNA-197, −196 and AAVS1, respectively (**Supplementary Table 5**). These DEGs are partially overlapping with those dysregulated in HD samples (from 43.0 to 52.3% for the up-regulated genes and from 27.5 to 40.7% for the down-regulated genes; **Figure 4C**). This suggests a higher susceptibility of SCD cells upon transfection and CRISPR/Cas9 treatment. As already observed for HD cells, most of the DEGs were dysregulated by the treatment with all sgRNAs (444 up- and 58 down-regulated genes; **Supplementary Figure 6B**).

**Figure 4.**
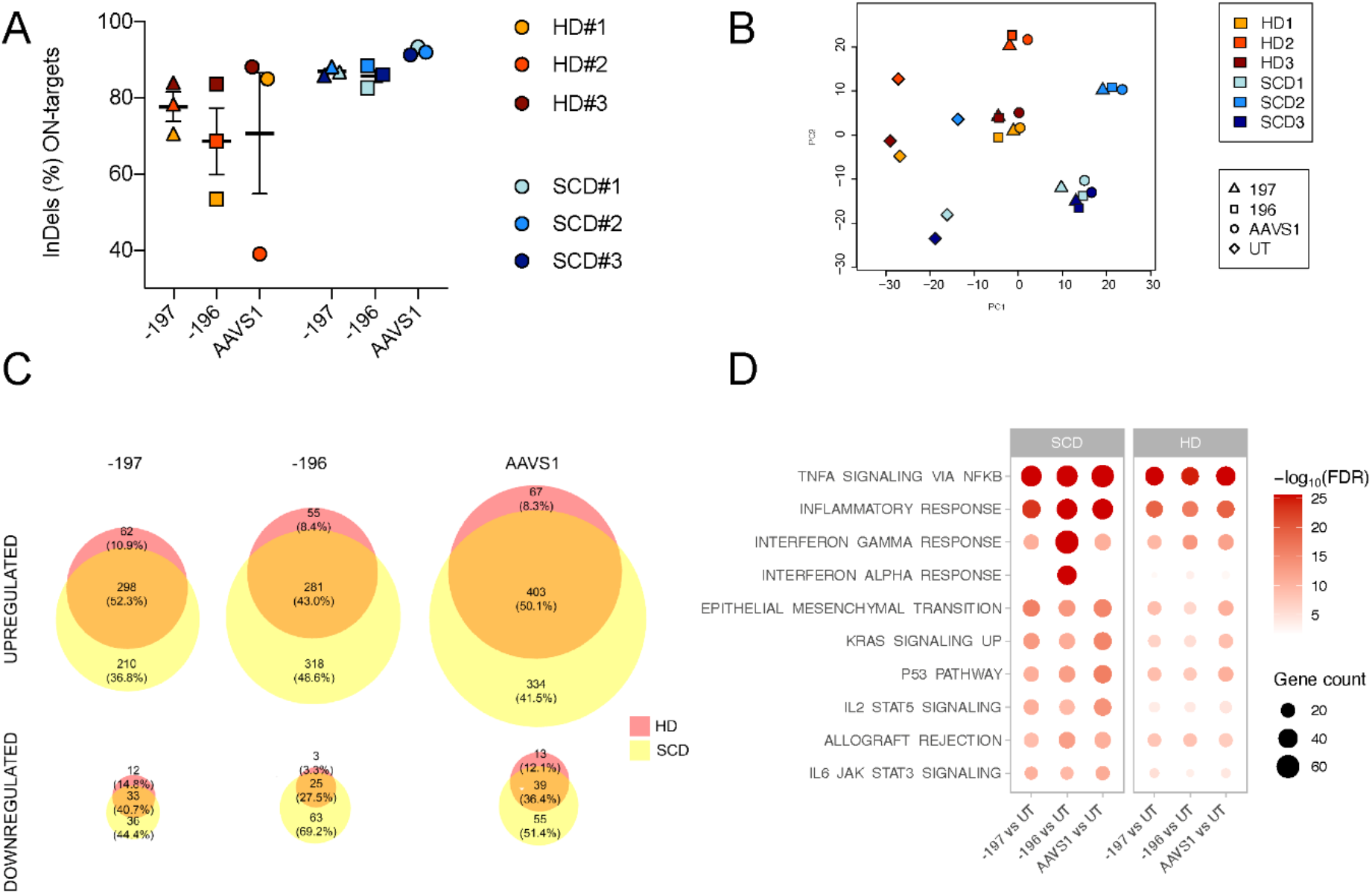
Transcriptomic changes following the genome editing procedure. (**A**) InDel frequency at *HBG1/2* promoters of Cas9-treated HD and SCD plerixafor-mobilized CD34^+^ cells, as evaluated by Sanger sequencing 6 days after transfection. (**B**) Unsupervised principal component analysis (PCA) of HD and SCD HSPCs two days after nucleofection. Different colors represent different donors. Shape symbols distinguish different sgRNA treatments (rhombuses: UT; triangles: −197, squares: −196; circles: AAVS1). (**C**) Venn diagrams showing the DEGs upregulated (upper panel) and downregulated (lower panel) upon treatment with the sgRNA −197, −196 and AAVS1 in HD (red) and SCD (yellow) treated HSPCs. (**D**) Functional enrichment analysis of up- regulated genes (FDR<0.05 and FC>2) in sgRNA-treated vs untreated (UT) samples from SCD patients and HD. The most enriched Hallmark gene sets are shown on the y-axis. The x-axis shows sample comparisons. The red color gradient indicates the statistical significance of the enrichment [expressed as –log10(FDR)]; the color scale values span from 0 to the 85th percentile of the dataset. The size of the circles reflects the quantity of genes associated with each Hallmark gene set.

We next performed a functional enrichment analysis focusing on MSigDB Hallmark gene sets to better investigate if genes up-regulated upon electroporation and CRISPR/Cas9 treatment are related to specific biological states or processes. Overall, these genes are involved in TNF-α signaling via NFKB, inflammatory response, interferon response, KRAS signaling, P53 pathway, allograft rejection and IL2/STAT5 and IL6/JAK/STAT3 signaling, and are typically activated upon electroporation and Cas9-mediated DSB formation^43^ (**Figure 4D and Supplementary Table 6**). Interestingly, we observed a stronger enrichment in SCD than in HD samples suggesting that HSPCs from SCD patients are more prone to inflammatory and P53 responses following electroporation and CRISPR/Cas9 treatment than HD cells (**Figure 4D**). Among the most up-regulated genes, we identified *CDKN1A*, a downstream effector of P53, known to be activated in response to DNA damage (**Supplementary Table 6)**. Enrichment in these same gene signatures was observed in all the samples regardless of the sgRNA or the genotype. However, it is worth noting a particular enrichment of genes involved in the interferon gamma and alpha response signature for the SCD samples treated with the sgRNA −196 (**Figure 4D** and **Supplementary Table 6**). In fact, these samples showed the strongest upregulation of several genes involved in the interferon-mediated response to RNA stimuli such as *OAS1*, *OAS2*, and *ISG15* (**Supplementary Table 6**).

## Discussion

We previously reported that Cas9-mediated disruption of the LRF BS in the *HBG1* and *HBG2* promoters induces a potent γ-globin reactivation, recapitulating the phenotype of asymptomatic SCD-HPFH patients^22^. Here, we first optimized the transfection procedure to achieve the highest editing efficiency while maintaining a good cell fitness. The use of a transfection enhancer and compounds maintaining stemness improved the editing protocol, while introducing a third NLS in the Cas9 construct failed to increase editing efficiency, suggesting that Cas9 nuclear localization is not a limiting factor. Interestingly, genome editing efficiency was modestly but consistently higher in SCD patient-derived HSPCs compared to HSPCs obtained from HDs, independently from the cell source (and therefore from the cell composition of different populations). These results suggest that SCD HSPCs are more prone to CRISPR/Cas9-mediated transfection and/or editing or might have differences in DNA repair mechanisms compared to HD HSPCs. This might be related to the high mutational burden observed in murine SCD HSPCs and the higher incidence of clonal hematopoiesis and leukemias reported in patients with SCD^38–40^.

Similarly, engrafted SCD cell populations showed increased levels of *HBG* targeting compared to HD HSCs in both primary and secondary recipients. This could be due to the highest editing efficiency in long-term repopulating HSCs from SCD patients. Alternatively, it could be partially ascribed to the highest editing efficiency observed in the input cell populations: while in mice transplanted with SCD HSPCs, all the input SCD HSCs are edited, in mice transplanted with HD HSPCs, unedited HSCs could have a competitive advantage compared to edited cells.

However, our data indicate a limited engrafting capability of SCD cells compared to HD HSPCs, even when using plerixafor-mobilized cells and even at steady state when transplanted untreated samples. Furthermore, for 3 out of 5 donors, electroporation of CRISPR/Cas9, further decreased (although modestly) the chimerism. Furthermore, while both untreated and edited SCD HSCs were able to differentiate *in vivo* in all the different blood lineages, we observed an increased differentiation of SCD cells towards CD14^+^ and CD11b^+^ myeloid cells in the BM of the recipient mice compared to HD HSCs. This finding was already reported by Park and colleagues^15^ when using non-mobilized SCD HSPCs and could be ascribed to the cell source. However, we observed increased frequencies of CD14^+^ and CD11b^+^ myeloid cells also when transplanting plerixafor-mobilized cells, suggesting a myeloid bias of HSCs potentially due the chronic inflammation in SCD cells^44^. Altogether, these results suggest that HSCs from SCD patients could be altered because of the inflammatory bone marrow microenvironment, and this phenotype could be exacerbated by the genome editing procedure.

Accordingly, our *in vitro* data (CFC assay) also showed an increased susceptibility to the electroporation and the CRISPR/Cas9 nuclease activity in SCD HSPCs. Furthermore, we evaluated the transcriptomic changes in matched HD and SCD plerixafor-mobilized HSPCs. Although, at steady state SCD and HD cells showed few dysregulated genes, the electroporation and the gene editing procedure induced a higher number of DEGs (involved in inflammatory and DNA damage responses) induced by the electroporation and the CRISPR/Cas9 activity^43^ in SCD samples compared to HD HSPCs. These results confirm the higher susceptibility of SCD cells and a particular cellular stress due to transfection and DSB formation. Importantly, most of the DEGs are commonly up- or down-regulated regardless of the editing site (*HBG* vs *AAVS1*), demonstrating that they are dysregulated by the transfection procedure and DSB formation rather than by the targeting of a specific locus. The only exception was the case on the −196 sgRNA that causes a stronger interferon response but only in SCD samples: whether this could be due to a particular cell stress, associated with the combination of the off-target activity or the inter-chromosomal translocation and the SCD “environment”, remains to be elucidated. Overall, this study highlights the need of performing safety studies, when possible, in clinically relevant conditions, i.e., in patient-derived HSPCs. Future experiments will require a more detailed analysis of *bona fide* HSCs (representing <1% of total HSPCs) at steady state and subjected to the genome editing procedure, e.g., single cell RNA-seq analyses.

Despite of these transcriptomic changes, our editing strategy still allows HSPC engraftment and is efficient in patient-derived HSCs: the transplantation of high doses of HSPCs in SCD patients - as currently performed in the CRISPR/Cas9-based clinical trial sponsored by Vertex^45^ using the FDA-approved Casgevy therapy - will likely circumvent the reduced engraftment capability of SCD HSCs.

The disruption of LRF BS resulted in efficient γ-globin reactivation in erythrocytes derived from engrafted human HSCs. Our data show that the percentage of HbF-expressing cells and the amount of γ-globin expression are positively correlated with the InDel frequency. If, as proposed^46,47^, HbF accounting for the 30% of the total hemoglobin is sufficient to ameliorate the clinical manifestations of SCD, our data indicate that this goal can be reached by disrupting three out of the four LRF BSs, which is likely, given the high editing efficiency (80%) achieved in SCD cells in our xenotransplantation experiments. These results are in line with previous studies based on the Cas9-targeting of γ-globin repressor binding sites at the *HBG* promoters^21^ and had prompted the researchers to maximize the InDel frequency of the HSPC input.

However, maximization of the editing efficiency is intrinsically linked to safety concerns associated to the unwanted DSBs and genomic rearrangements^24–30^. This is particularly relevant in the context of SCD, where patients’ HSPCs might be characterized by a high mutational burden. In our study, CAST-seq (performed in primary HSPCs) identified two off-target sites (one for each sgRNA) with high InDel frequencies (up to ∼40% in SCD cells *in vitro* and *in vivo*). Importantly, one of them was not described in our previous work^22^ because of the limitations of GUIDE-seq, a method that identify off-target in a cell line, which could not reflect the chromatin structure of primary cells. Furthermore, while GUIDE-seq identified many off-targets that were not validated by targeted amplicon sequencing in primary cells^22^, the main off-targets identified by CAST-seq were confirmed by sequencing analyses. Therefore, as CAST-seq in primary HSPCs and GUIDE-seq in 293T cells can identify different off-targets, they might represent complementary analyses to monitor the safety of genome editing approaches.

Off-target editing also increases the risk for chromosomal rearrangements. In this study, we reported the occurrence of an inter-chromosomal translocation between on- and off-target sites in samples edited with the −196 sgRNA. The frequency of these events was low but seemed to be more pronounced in SCD cells *in vitro*, again suggesting a correlation with the potentially higher mutational burden in SCD compared to HD HSPCs. Given the need of high DNA amount, we could not attempt to detect this translocation *in vivo* to rule out if HSPCs with large genomic rearrangements are counter-selected during the engraftment or differentiation.

Finally, as previously described^21^, we reported the frequent occurrence both *in vitro* and *in vivo* of 4.9-kb deletions caused by the simultaneous cleavage of the identical *HBG1* and *HBG2* promoters. These events led to loss of the *HBG2* gene and ^G^γ-globin expression.

Importantly both off-target and intra- or inter-chromosomal rearrangements appear to be well tolerated by HSPCs as they do not alter target gene expression (except for a reduced ^G^γ-globin expression) and do not impair the engrafting and differentiation ability of HSPCs. However, we cannot exclude that these unintended events can have long-term, hardly predictable toxic effects.

While given the intrinsic nature of the target (nearly identical paralog *HBG1* and *HBG2* genes) it is difficult to avoid the generation of the 4.9-kb deletions, high fidelity Cas9 could be used to decrease off-targets (and therefore inter-chromosomal translocations). These variants have proved their efficacy in reducing but not abolishing off-target activity, while often affecting on-target efficiency^48^. It is noteworthy that a lower on-target editing (and as a consequence a reduced off-target activity) achieved either by decreasing the exposure to the CRISPR/Cas9 (e.g., by reducing the RNP amount) or by using high-fidelity Cas9 could be envisaged, given the selective advantage of corrected RBCs in SCD patients^47,49^, as shown for the Casgevy gene therapy^45^.

More in general, the use of nucleases and the consequent DSB formation can be associated with harmful side effects that significantly limit their safety profile. For this reason, recent approaches based on DSB-free technologies have been developed to reactivate HbF (by reproducing HPFH mutations^50–52^ or by downregulating the expression of BCL11A, a major HbF transcriptional repressor^53^) or to revert the SCD causing mutation^54–56^. By way of example base and prime editing strategies targeting the *HBG1/2* promoters can substantially reduce the generation of the 4.9 kb deletions^52^. However, despite the promise of reduced off-target activities, prime editing strategies are still relatively inefficient in primary SCD cells^56^. Furthermore, while highly efficient in *bona fide* HSCs, base editors show both DNA and RNA off-target activities that need to be carefully evaluated before moving to the clinic in relevant HSPCs from SCD patients, given their unique properties compared to normal HSPCs identified in our study.

## Materials and Methods

### HSPC purification and culture

Peripheral blood human plerixafor or granulocyte colony-stimulating factor (G-CSF)-mobilized adult HSPCs or cord blood HSPCs were obtained from HDs. Peripheral blood non-mobilized or plerixafor-mobilized human adult HSPCs were obtained from SCD patients. SCD and HD samples eligible for research purposes were obtained from the Necker-Enfants malades Hospital (Paris, France) except plerixafor-mobilized HD cells that were purchased from Caltag and Hemacare. Written informed consent was obtained from all the subjects. All experiments were performed in accordance with the Declaration of Helsinki. The study was approved by the regional investigational review board (reference: DC 2022-5364, CPP Ile-de-France II “Hôpital Necker-Enfants malades”). HSPCs were purified by immunomagnetic selection after immunostaining using the CD34 MicroBead Kit (Miltenyi Biotec). HSPCs were thawed and cultured at a concentration of 5×10^5^ cells/ml in the “HSPC medium” containing StemSpan (STEMCELL Technologies) supplemented with penicillin/streptomycin (Gibco), 250 nM StemRegenin1 (STEMCELL Technologies), 20 nM UM171 (STEMCELL Technologies), and the following recombinant human cytokines (PeproTech): human stem cell factor (SCF; 300 ng/ml), Flt-3L (300 ng/ml), thrombopoietin (TPO; 100 ng/ml), and interleukin-3 (IL-3; 60 ng/ml).

### Ribonucleoprotein (RNP) transfection

RNP complexes were assembled at room temperature using a Cas9:sgRNA ratio of 1:2 [90 pmol of the Cas9 (Cas9x2NLS_GFP, Cas9x2NLS or Cas9x3NLS) protein and 180 pmol of the synthetic sgRNA; Synthego]. HSPCs (2×10^5^ cells/condition) were transfected with RNP complexes using the P3 Primary Cell 4D-Nucleofector X Kit (Lonza) and the CA137 program (Nucleofector 4D) with or without a transfection enhancer (IDT). Untreated cells or cells transfected with RNP complexes containing a sgRNA targeting the *AAVS1* locus, served as negative controls^22^.

### CFC-assay

HSPCs were plated at a concentration of 1 × 10^3^ cells/ml in a methylcellulose-containing medium (GFH4435, STEMCELL Technologies) under conditions supporting erythroid and granulo-monocytic differentiation. BFU-E and CFU-GM colonies were scored after 14 days. BFU-Es and CFU-GMs were randomly picked and collected as bulk populations (containing at least 25 colonies) or as individual colonies (35 to 45 colonies per sample) to evaluate genome editing efficiency and/or globin expression.

### Evaluation of InDel frequency and large genomic rearrangements

InDel frequency and the presence of genomic rearrangements were evaluated in HSPCs 6 days after transfection, in BFU-Es and CFU-GMs 14 days after plating, in human CD45^+^ cells sorted from the BM of NSG recipient mice, and in spleen derived from the same mice. Genomic DNA was extracted from control and edited cells using the PURE LINK Genomic DNA Mini kit (LifeTechnologies) following manufacturer’s instructions.

#### InDel frequency

The InDel frequency was evaluated at the on- or off-target sites by PCR, using the primers listed in **Table 1**, followed by Sanger sequencing and TIDE analysis (Tracking of InDels by Decomposition)^50^.

**Table 1.**
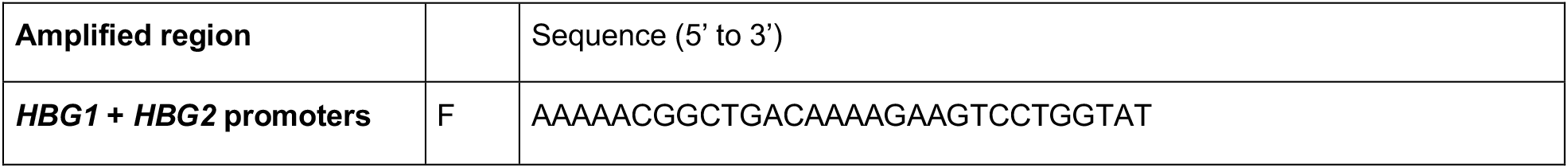
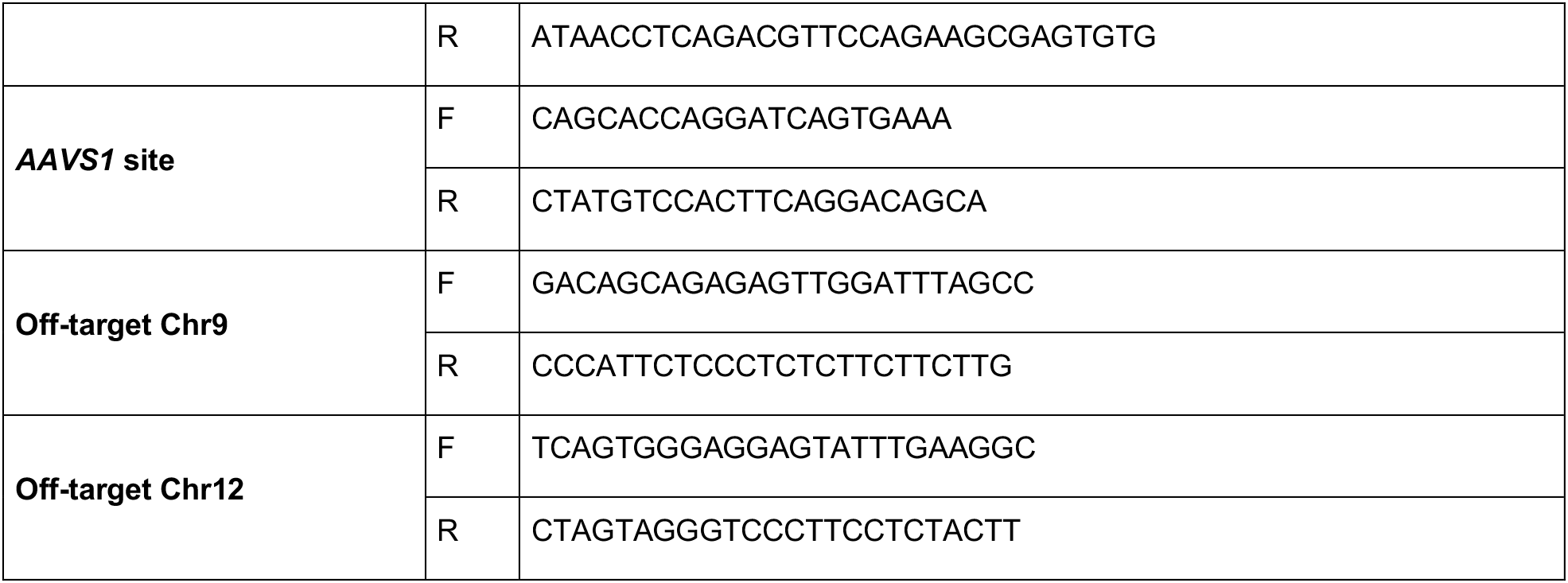
Primers used to detect InDels events. F, forward primer; R, reverse primer.

#### Detection of the 4.9-kb deletion and inter-chromosomal translocation by ddPCR

Digital Droplet PCR (ddPCR) were performed using primers and probes listed in **Table 2** to quantify the frequency of the 4.9-kb deletion and the inter-chromosomal translocation between *HBG* and the off-target located on chromosome 12. PCR using control primers annealing to *hALB* (located on chromosome 4) and *hRAD1* (located on chromosome 5) were used as DNA loading control for 4.9-kb deletion and inter-chromosomal translocation, respectively.

**Table 2.**
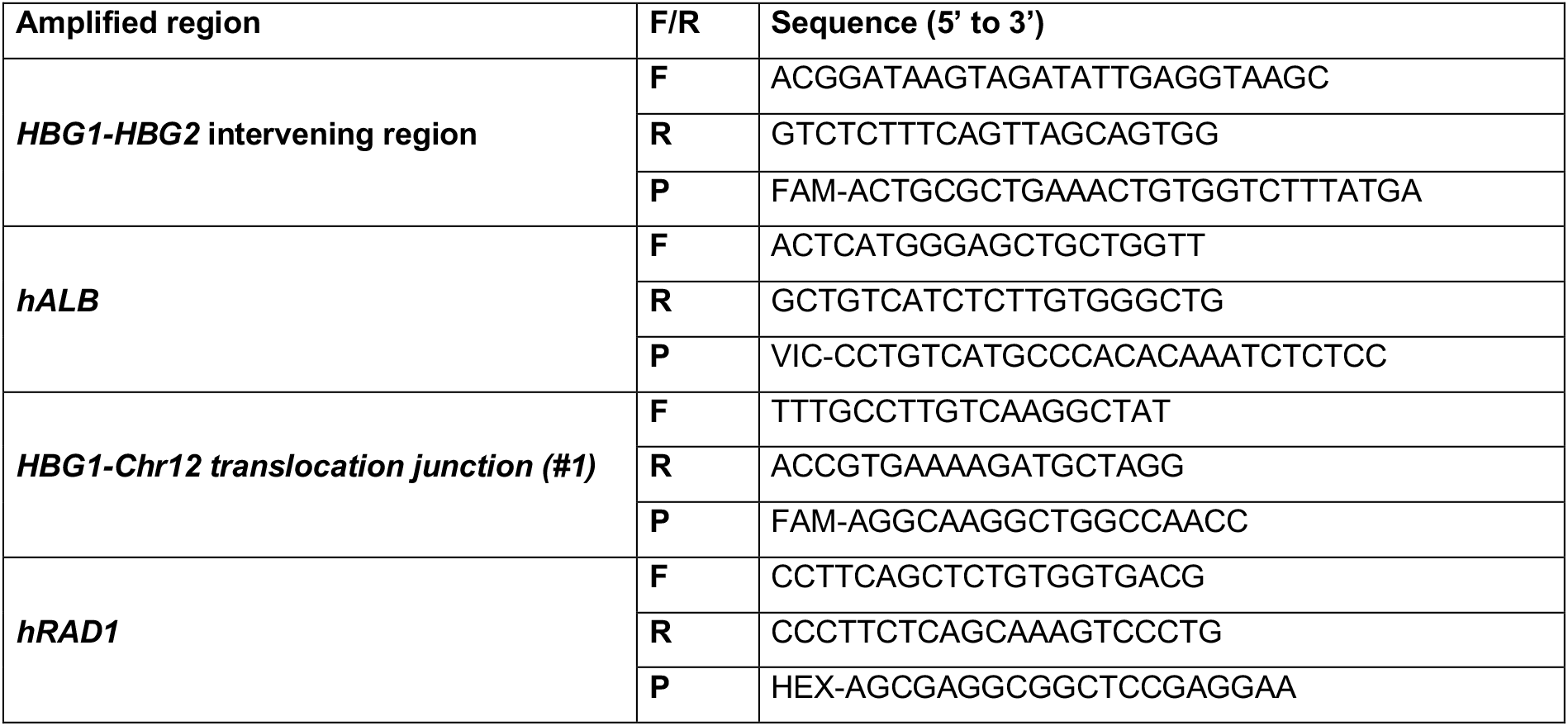
Primers used for ddPCR. F, forward primer; R, reverse primer; P, probe.

#### CAST-seq

Chromosomal Aberration analysis by Single Targeted Linker Mediated PCR method (CAST-seq) was performed as described in^30^ with the following modifications. Genomic DNA, extracted from edited and mock treated cells, was fragmented, repaired, and A-tailed using the NEB Next Ultra II FS DNA Library Prep Kit for Illumina (New England Biolabs). Linkers were then ligated to the DNA using the NEB Next Ultra II Ligation Master Mix and Ligation Enhancer (New England Biolabs). The CAST-seq PCR reactions were performed in forward and in reverse in respect to the cleavage site. We used the primers listed in **Table 3**.

**Table 3.**
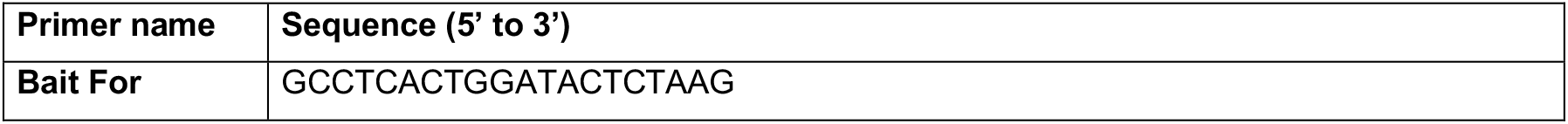
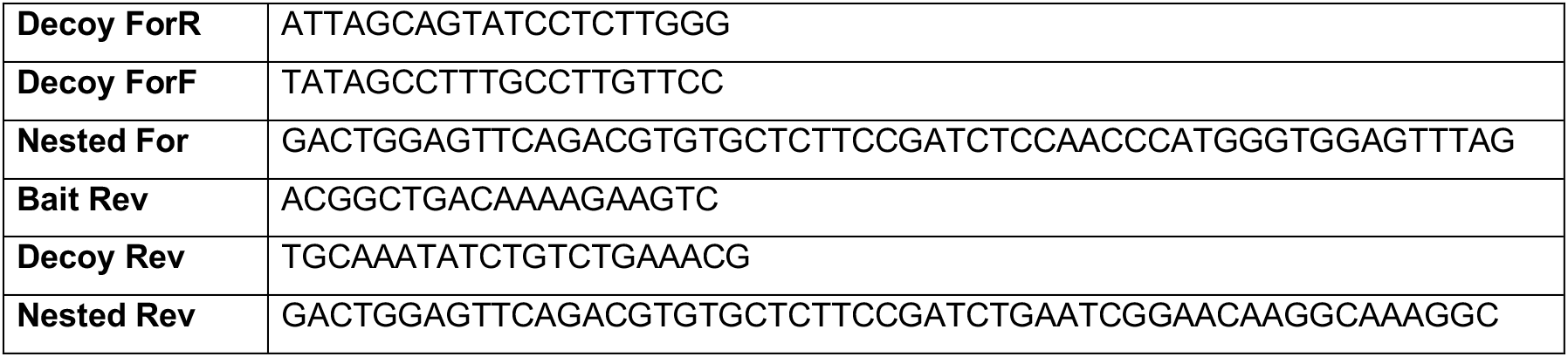
Primers used to detect Cas9-mediated translocation by CAST-seq.

The reactions were purified, barcoded with NEB Next Multiplex Oligos for Illumina (New England Biolabs) and sequenced by Illumina MiSeq (MiSeq Reagent Kit v3 600-cycle). Bioinformatic analysis of the sequences was performed as previously described^30^ identifying true translocation sites when those events are present in two replicates and significant when compared to the untreated samples (p < 0.05).

#### Inter-chromosomal translocations

In order to detect inter-chromosomal translocations, we perform a PCR using primers designed across on-target (*HBG*) and off-target sites identified by CAST-seq. 200 ng of genomic DNA derived from treated HSPCs was used with different combinations of Fw and Rev primers (listed in **Table 4**). Amplicons were cloned into pCR 2.1-TOPO TA vector (ThermoFischer), and then transformed into One Shot™ TOP10 Chemically Competent (ThermoFischer) following manufacturer’s instructions. Plasmids purified from 10 bacterial colonies were subjected to Sanger sequencing.

**Table 4.**
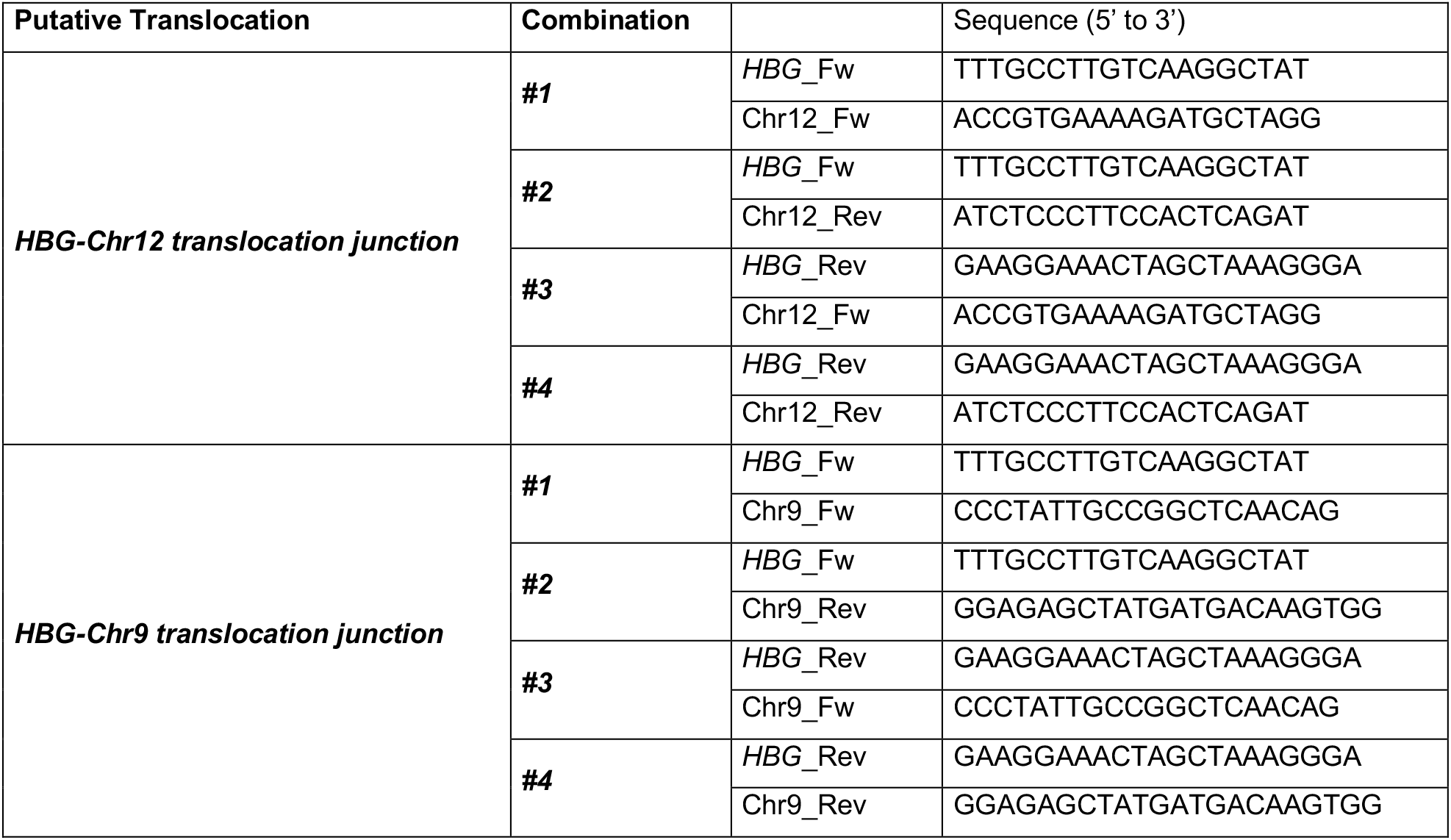
Primers used for regular PCR for the identification of inter-chromosomal translocation.

Digital Droplet PCR (ddPCR) was performed using primers and probes listed in **Table 2** to quantify the frequency of chromosomal translocations.

### RP-HPLC analysis of globin chains

Reverse-phase (RP)-HPLC analysis was performed using a NexeraX2 SIL-30AC chromatograph and the LC Solution software (Shimadzu). A 250x4.6 mm, 3.6 μm Aeris Widepore column (Phenomenex) was used to separate globin chains by RP-HPLC. Samples were eluted with a gradient mixture of solution A (water/acetonitrile/trifluoroacetic acid, 95:5:0.1) and solution B (water/acetonitrile/trifluoroacetic acid, 5:95:0.1). The absorbance was measured at 220 nm.

### CE-HPLC analysis of hemoglobin tetramers

Cation-exchange HPLC analysis was performed using a NexeraX2 SIL-30AC chromatograph and the LC Solution software (Shimadzu). A 2 cation-exchange column (PolyCAT A, PolyLC, Columbia, MD) was used to separate hemoglobin tetramers by HPLC. Samples were eluted with a gradient mixture of solution A (20mM bis Tris, 2mM KCN, pH=6.5) and solution B (20mM bis Tris, 2mM KCN, 250mM NaCl, pH=6.8). The absorbance was measured at 415 nm.

### RT-qPCR

Total RNA was extracted from SCD or HD HSPCs (15, 24, 48, 72 hours and 6 days post-transfection) using the Quick-DNA/RNA Miniprep (ZYMO Research). RNA was treated with DNase using the DNase I kit (Invitrogen) following manufacturer’s instructions. Mature transcripts were reverse-transcribed using the SuperScript First-Strand Synthesis System for RT-qPCR (Invitrogen) with oligo (dT) primers. RT-qPCR was performed using the iTaq universal SYBR Green master mix (Biorad), the primers listed in **Table 5**, and the Viia7 Real-Time PCR system (ThermoFisher Scientific), or the CFX384 Touch Real-Time PCR Detection System (Biorad).

**Table 5.**
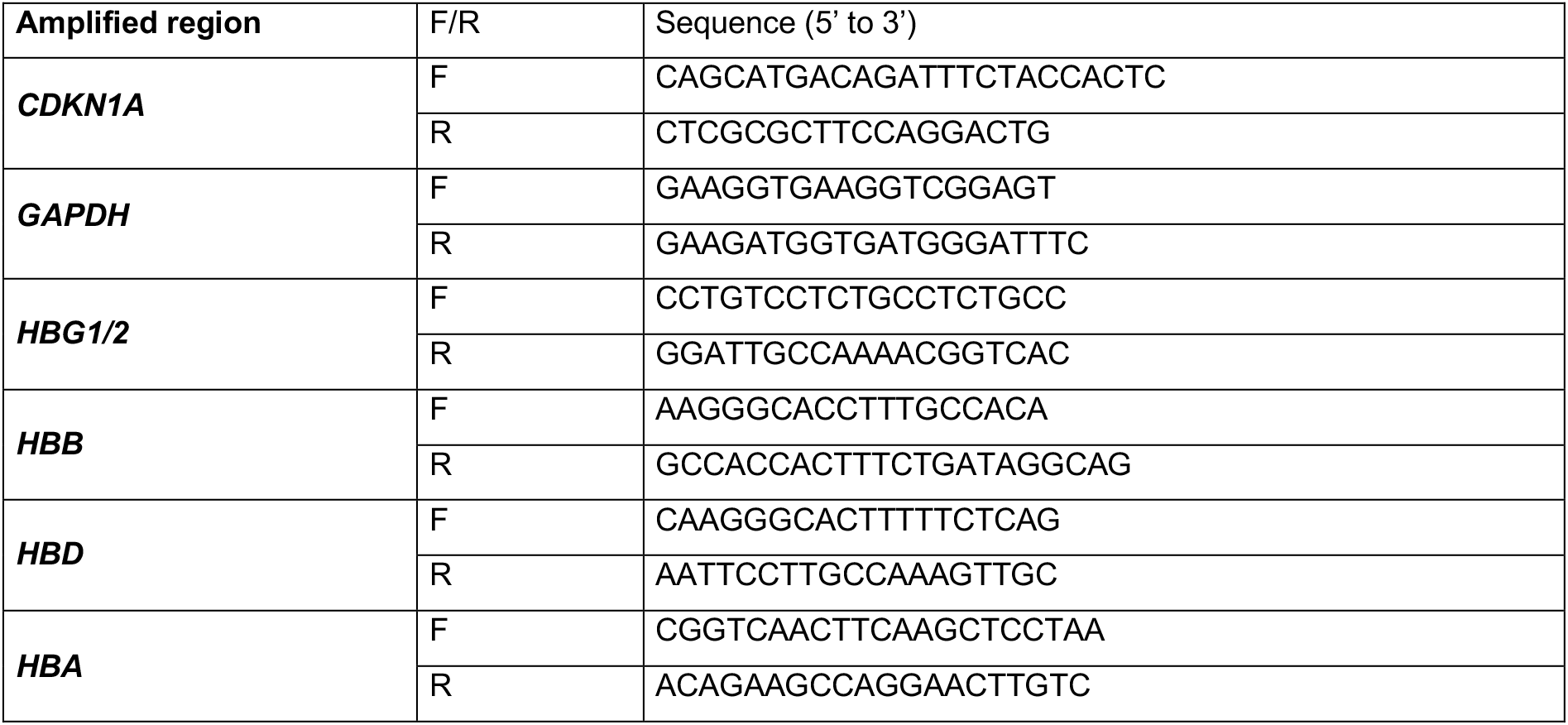
Primers used for RT-qPCR. F, forward primer; R, reverse primer.

### Flow cytometry analysis

Transfected HSPCs were characterized for the expression of CD34, CD38, CD133, and CD90, using an APC-Cy7- conjugated anti-CD34 antibody (343514, Biolegend), a PE-Cy7-conjugated anti-CD38 antibody (555462, BD Pharmingen), a PE-conjugated anti-CD133 antibody (130-113-748, Miltenyi) and an APC-conjugated anti-CD90 antibody (555596, BD Pharmingen).

Cells harvested from femurs and spleen of the host mice were stained with antibodies against murine or human surface markers [vioblue-conjugated anti-murine CD45 antibody (130-110-664, Miltenyi), APC-Vio770- conjugated anti-human CD45 antibody (130-110-635, Miltenyi), APC-conjugated anti-human CD3 antibody (130-113-135, Miltenyi), PE-Cy7-conjugated anti-human CD14 (562698, BD Pharmingen), PE-conjugated anti- human CD15 (130- 113-485, Miltenyi), BV510-conjugated anti-human CD19 (562947, BD Pharmingen), PE-conjugated anti-human CD235a/GYPA (555570, BD Pharmingen), APC-conjugated anti-human CD11b (553312, BD Pharmingen), PE-Vio 770-conjugated anti-human CD34 (130-124-456, BD Miltenyi) and FITC-conjugated anti-human CD36 (555454, BD Pharmingen)].

Erythroid cells were fixed with 0.05% cold glutaraldehyde and permeabilized with 0.1% TRITON X-100. After fixation and permeabilization, cells were stained with a PE-Cy7-conjugated antibody recognizing the CD235a/GYPA erythroid surface marker (563666, BD Pharmingen) and a FITC-conjugated antibody recognizing HbF (clone 2D12 552829, BD). Flow cytometry analysis of CD36, CD71, CD235a/GYPA, BAND3 and α4-Integrin erythroid surface markers was performed using a V450-conjugated anti-CD36 antibody (561535, BD Horizon), a FITC-conjugated anti-CD71 antibody (555536, BD Pharmingen), a PE-Cy7-conjugated anti-CD235a/GYPA antibody (563666, BD Pharmingen), a PE-conjugated anti-BAND3 antibody (9439, IBGRL) and an APC-conjugated anti-CD49d antibody (559881, BD). Flow cytometry analysis of enucleated or viable cells was performed using double-stranded DNA dyes (DRAQ5, 65-0880-96, Invitrogen and 7AAD, 559925, BD, respectively).

Flow cytometry analyses were performed using Fortessa X20 (BD Biosciences) or Gallios (Beckman Coulter) flow cytometers. Data were analyzed using the FlowJo (BD Biosciences) software.

### Xenotransplantation

Nonobese diabetic severe combined immunodeficiency gamma (NSG) mice (NOD.CgPrkdcscid Il2rgtm1Wj/SzJ, Charles River Laboratories, St Germain sur l’Arbresle, France) were housed in a specific pathogen-free facility. For primary and secondary transplantation, mice at 6 to 8 weeks of age were conditioned with busulfan (Sigma, St. Louis, MO, USA) injected intraperitoneally (25 mg/kg body weight/day) 24, 48, and 72 hours before transplantation. Neomycin and acid water were added in the water bottle. For primary transplantation, control or edited mobilized (0.5-1.0 x 10^6^ cells per mouse) or non-mobilized (0.5-2.0 x 10^6^ cells per mouse) HSPCs from HD or SCD patients were transplanted into NSG mice via retro-orbital sinus injection. For secondary transplantation, half of the BM from one or two primary recipient mice were transplanted into NSG mice via retro-orbital sinus injection. At 16 weeks after transplantation, NSG mice were sacrificed. All experiments and procedures were performed in compliance with the French Ministry of Agriculture’s regulations on animal experiments and were approved by the regional Animal Care and Use Committee (APAFIS#2101-2015090411495178 v4).

### *Ex vivo* erythroid differentiation of human CD45^+^ cells

Human CD45^+^ cells were differentiated into mature RBCs using a three-phase erythroid differentiation protocol^51,52^. During the first phase (day 0 to day 6), cells were cultured in a basal erythroid medium supplemented with 100 ng/ml recombinant human SCF (PeproTech), 5 ng/ml recombinant human IL-3 (PeproTech), 3 IU/ml EPO Eprex (Janssen-Cilag) and 10^−6^ M hydrocortisone (Sigma). During the second phase (day 6 to day 9), cells were co-cultured with MS-5 stromal cells in the basal erythroid medium supplemented with 3 IU/ml EPO Eprex (Janssen-Cilag). During the third phase (day 9 to day 20), cells were co-cultured with stromal MS-5 cells in a basal erythroid medium without cytokines. From day 13 to 20, human AB serum was added to the medium. Erythroid differentiation was monitored by flow cytometry analysis of CD36, CD71, CD235a/GYPA, BAND3 and α4-Integrin erythroid surface markers and of enucleated cells using the DRAQ5 double-stranded DNA dye. 7AAD was used to identify live cells.

### RNA-seq

Total RNA was isolated from untreated or RNP-transfected plerixafor-mobilized HD and SCD HSPCs (n=3 for each group) using the RNeasy Kit (QIAGEN) that includes a DNAse treatment step. RNA quality was assessed by capillary electrophoresis using High Sensitivity RNA reagents with the Fragment Analyzer (Agilent Technologies) and the RNA concentration was measured using both Xpose spectrophotometry (Trinean) and Fragment Analyzer (Agilent Technologies) capillary electrophoresis.

RNA-seq libraries were prepared starting from 30 ng of total RNA using the Universal Plus mRNA-Seq kit (Nugen) as recommended by the manufacturer. Briefly, mRNA was captured with polyA+ magnetic beads from total RNA. mRNA was chemically fragmented. Single strand and second strand cDNA were produced and then ligated to Illumina compatible adaptors with UDI. To produce oriented RNA-seq libraries, a final step of strand selection was performed. The NuQuant system (Nugen) was used to quantify the RNA-seq libraries. An equimolar pool of the final indexed RNA-Seq libraries was prepared and sequenced using the Illumina NovaSeq 6000 system (paired-end sequencing; 2×100-bp). A total of ∼50 millions of passing filter paired-end reads were produced per library.

Read quality was verified using FastQC (v. 0.11.9^57^). Raw reads were trimmed for adapters and low-quality tails (quality < Q20) with BBDuk (v. 38.92^58^); moreover, the first 15 nucleotides were force-trimmed for low quality. Reads shorter than 35 bp after trimming were removed. Reads were subsequently aligned to the human reference genome (hg38) using STAR (v. 2.7.9a^59^). Raw gene counts were obtained in R-4.1.1 using the *featureCounts* function of the *Rsubread* R package (v. 2.6.4^60^) and the GENCODE 38 basic gene annotation for hg38 reference genome. Gene counts were normalized to counts per million mapped reads (CPM) and to fragments per kilobase of exon per million mapped reads (FPKM) using the *edgeR* R package (v. 3.34.1^61^); only genes with a CPM greater than 1 in at least 3 samples were retained for differential analysis. Differential gene expression analysis was performed using the *glmQLFTest* function of the *edgeR* R package, using donor as a blocking variable.

Functional enrichment analysis on MsigDB Hallmark gene sets was performed using the ToppFun tool of the ToppGene suite^62^.

## Data availability statement

The datasets supporting the results of this article are available in the Gene Expression Omnibus repository under the accession number GSE248671.

## Acknowledgments

This work was supported by state funding from the French National Research Agency (*Agence Nationale de la Recherche*) as part of the *Investissements d’Avenir* program (ANR-10-IAHU-01), the Paris Ile de France Region (under the “DIM Thérapie génique” initiative), the European Research Council (865797 DITSB), the European Commission (HORIZON-RIA EDITSCD grant no. 101057659) and the AFM-Telethon (grants 22206, 22399 and 23879).

## Author Contributions

GF designed, conducted experiments, analyzed data and wrote the paper, MB analyzed data and wrote the paper, GS, BM, TF, AC, PA, GH and GT conducted experiments and analyzed data. JPC contributed to the design of the experimental strategy. OR analyzed RNA-seq data. AM conceived the study, designed experiments and wrote the paper.

---

**Supplementary Figure 1.**
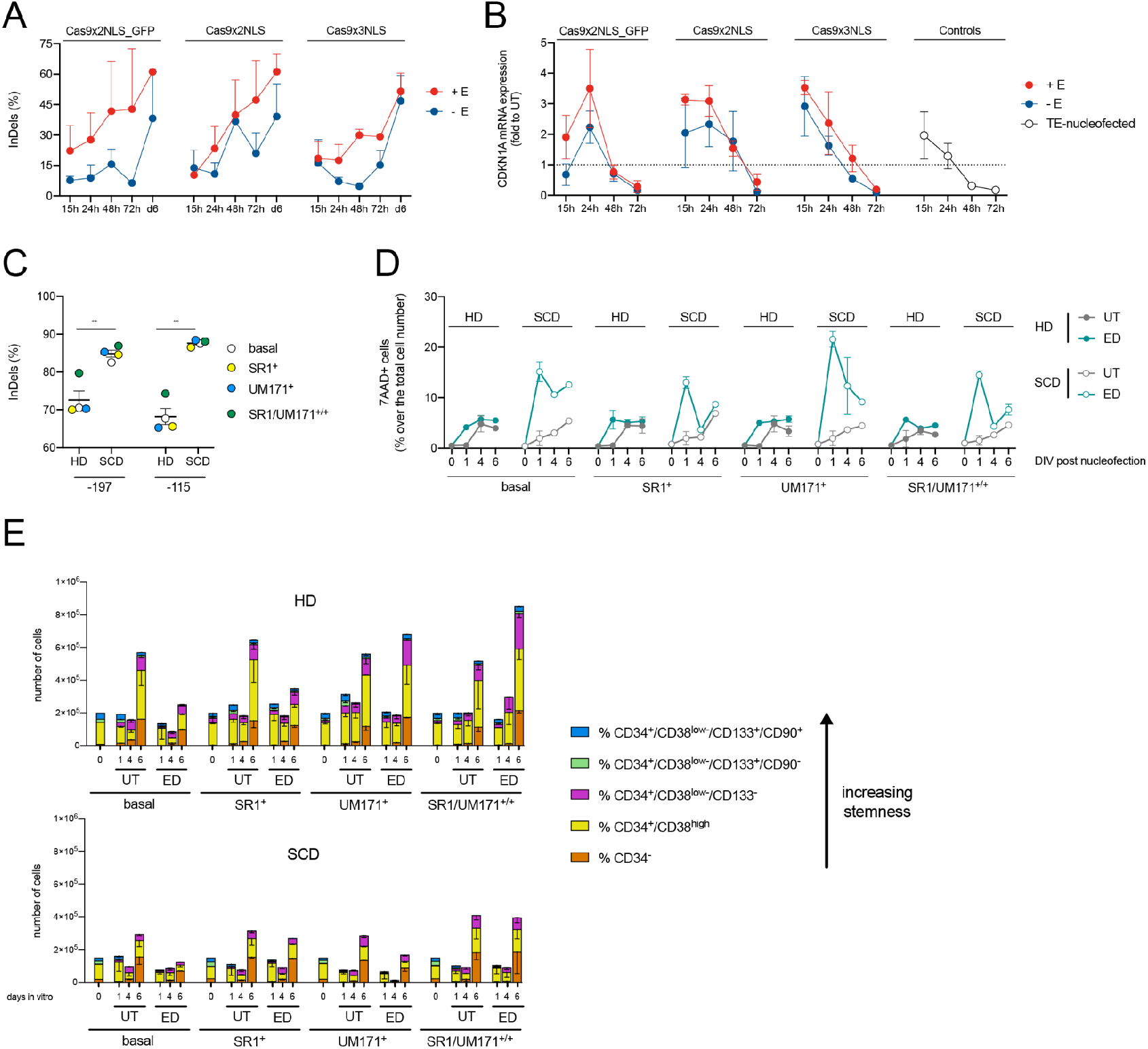
Optimization of the genome editing procedure in primary HSPCs. (**A**) Time course analysis of InDel frequency measured in CB-derived HD HSPCs treated with three different Cas9 RNPs (Cas9x2NLS_GFP, Cas9x2NLS, Cas9x3NLS) in the presence (+E, red line) or in the absence (-E, blue line) of the electroporation enhancer. We used the −197 sgRNA. (**B**) Time course analysis of *CDKN1A* mRNA expression in CB-derived HD HSPCs treated with three different Cas9 RNP nucleases (Cas9x2NLS_GFP, Cas9x2NLS, Cas9x3NLS) in the presence (+E, red line) or in the absence (-E, blue line) of the electroporation enhancer. Data are reported as mean ± SEM of n=2 run in triplicate. **p<0.01; paired t-test. (**C**) GE efficiency in cord blood-derived (HD) and non-mobilized (SCD) CD34+ cells measured by Sanger sequencing followed by TIDE analysis in samples edited with two different *HBG*-targeting sgRNA (the −197 sgRNA or the −115 sgRNA targeting the −115 region of the *HBG* promoters^2^). Each colored dot indicates a different culture condition. Data are reported as mean ± SEM of 4 replicates. (**D**) Time course analysis of the percentage of dead cells (measured as 7AAD^+^ cells) in edited (ED, green line) and control samples (UT, grey line) in HD- (filled dots) vs SCD (empty dots) patient-derived HSPCs cultured in the presence (+) or in the absence (-) of SR1 and/or UM171. Data are reported as mean ± SEM of 2 replicates. (**E**) Bar plots showing the cell composition of HD- (top panel) and SCD (bottom panel) patient-derived HSPCs cultured in the presence (+) or in the absence (-) of SR1 and/or UM171. We defined 5 populations with increasing stemness properties (CD34- cells, CD34+/CD38high cells, CD34+/CD38low-/CD133- cells, CD34+/CD38low-/CD133+/CD90- cells, and CD34+/CD38low-/CD133+/CD90+ cells). Starting from day 4, compared to HD samples, SCD cells had a larger fraction of more differentiated CD34^-^ cells and the CD34^+^/CD38low^-^/CD133^+^/CD90^+^ cell population (more enriched in HSCs) disappeared in SCD samples. In fact, this latter fraction is still present in HD cells at day 4, although it is smaller compared to the beginning of the culture. However, it is worth to notice that since the beginning SCD samples have a higher proportion of CD34^-^ cells and a lower percentage of CD34^+^/CD38low^-^/CD133^+^/CD90^+^ cells, which could be ascribed to the cell origin (patient adult vs HD cord blood HSPCs). Data are reported as mean ± SEM of 2 replicates.

**Supplementary Figure 2.**
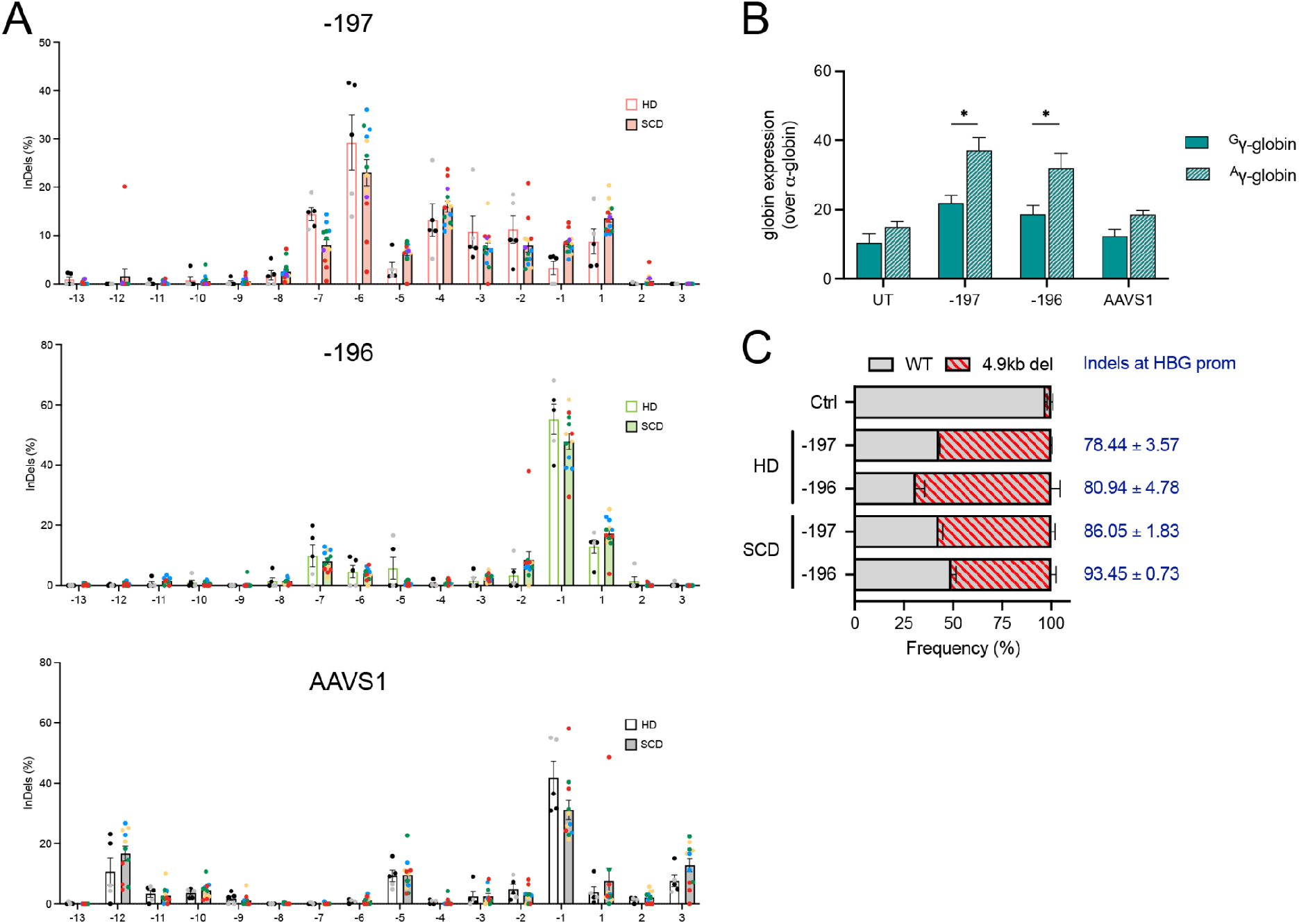
Editing profiles in primary SCD and HD HSPCs. (**A**) Single frequency of individual InDels in healthy donor (HD) and sickle cell disease (SCD) HSPCs treated with the −197, −196 and AAVS1 sgRNAs. InDel frequency was evaluated by Sanger sequencing followed by TIDE analysis. n= 2 donors for HD; n=5 donors for SCD. (**B**) Expression of ^G^γ- and ^A^ γ-globin chains measured by RP-HPLC in pooled SCD BFU-E colonies derived from untreated (UT) HSPCs and HSPCs treated with the −197, −196 and AAVS1 sgRNA. Data are expressed as mean ± SEM n=4, *p<0.05 Mann-Whitney test. (**C**) Frequency of 4.9-kb deletion as measured by ddPCR in healthy donor (HD) and sickle cell disease (SCD) HSPCs. We also reported mean ± SEM values of the overall InDels measured by Sanger sequencing and TIDE analysis at the *HBG1/2* promoters. UT and AAVS1-treated cells served as negative control (Ctrl).

**Supplementary Figure 3.**
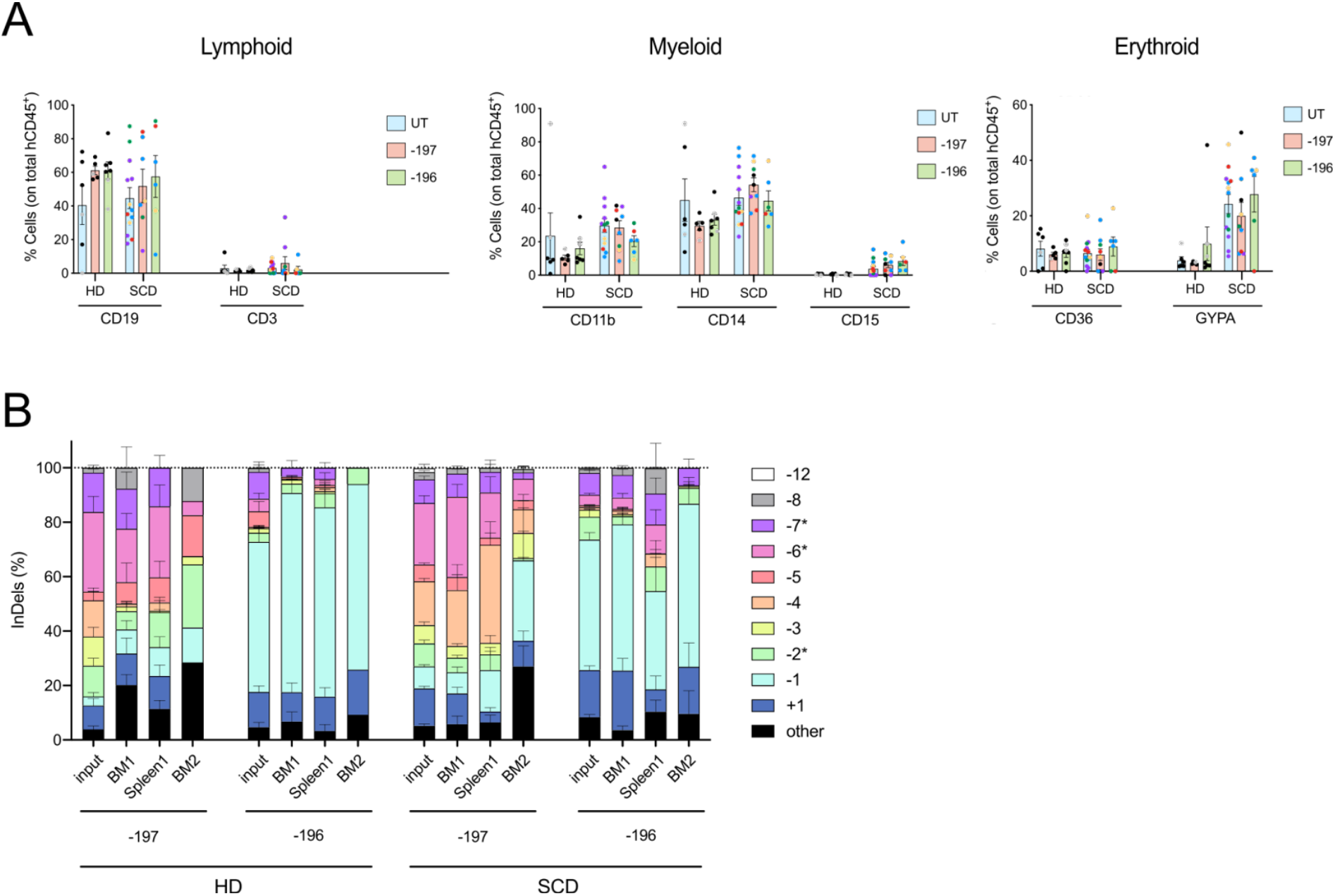
Multilineage differentiation and editing profiles in primary SCD and HD HSPCs. (**A**) Frequency of human T (CD3) and B (CD19) lymphoid, myeloid (CD14, CD15 and CD11b) and erythroid (CD36, GYPA) cells in the spleen of mice transplanted with control and edited HSPCs. (**B**) Frequency (expressed as %) of individual InDel events in the input populations and in BM- and spleen-derived human CD45+ cells edited with the −197 and −196 sgRNAs, as evaluated by Sanger sequencing and TIDE analysis. The numbers 1 and 2 after the organs’ name indicate primary and secondary recipients, respectively. * indicate putative microhomology associated events.

**Supplementary Figure 4.**
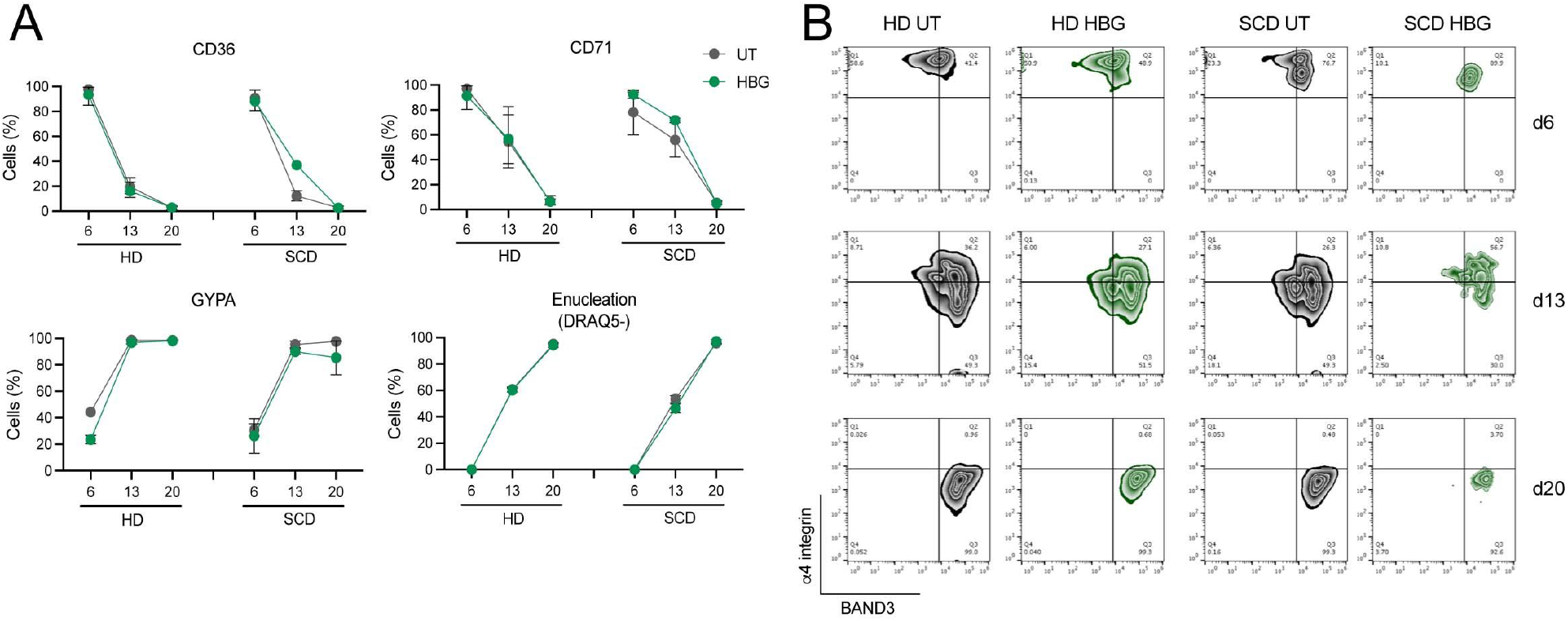
Erythroid differentiation of edited HSPCs. (**A**) Flow cytometry analysis of early (CD36 and CD71) and late (Glycophorin A GYPA) erythroid markers, and enucleation rate (measured as frequency of DRAQ5^-^ cells) in *ex vivo* differentiated erythrocytes. Data are plotted as means ± SEM (n=2-6). (**B**) Representative flow-cytometry plots showing the expression of BAND3 and α4 integrin in 7AAD^-^ GYPA^+^ cells at day (d) 6, 13 and 20 of *ex vivo* erythroid differentiation of the BM-repopulating hCD45^+^ cells. Edited samples (HBG) are highlighted in green and untreated (UT) samples in grey.

**Supplementary Figure 5.**
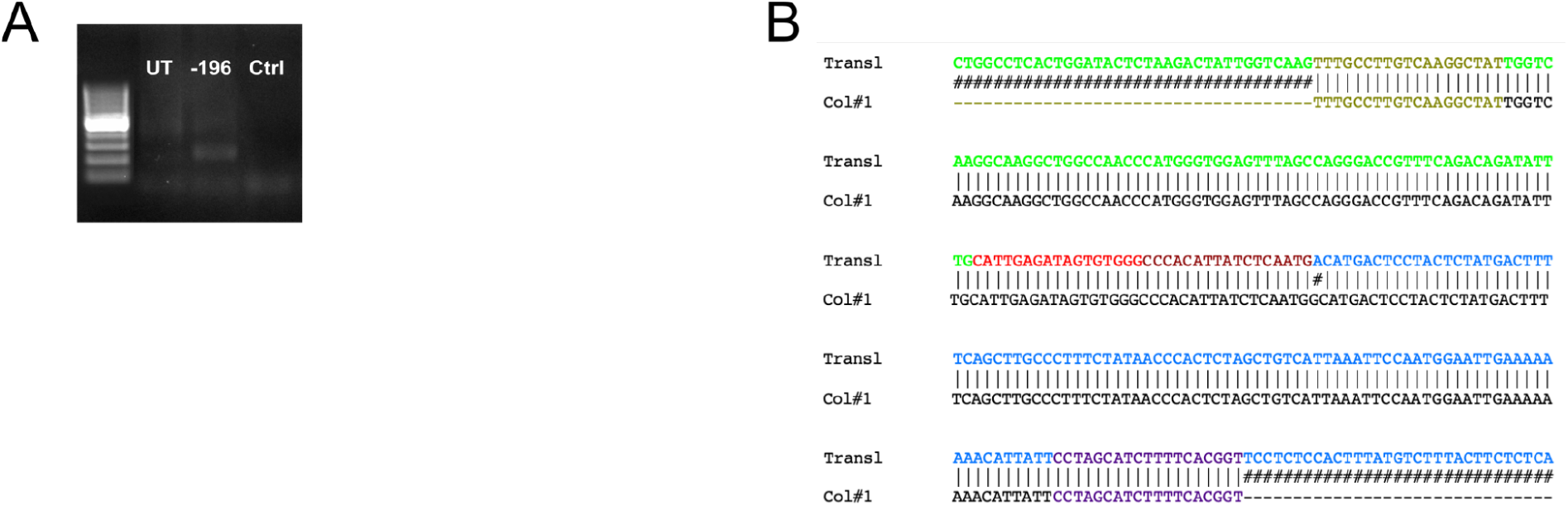
Inter-chromosomal translocation in HSPCs treated with the sgRNA −196. (**A**) Agarose gel image showing a band corresponding to the PCR product obtained using primers surrounding the putative translocation junctions in untreated (UT) SCD HSPCs or SCD HSPCs treated with the sgRNA −196 (−196). As control, we performed the PCR reaction in the absence of DNA (ctrl). (**B**) Alignment between the *in silico* predicted translocation (Trans1) and the sequence of the PCR product showed in A (Col#1). Green and blue identify the *HBG* promoter and off-target on chromosome 12 sequence, respectively. We indicated in red and brown the sequences of the −196 sgRNA on-target site and of the off-target site on chromosome 12 (without the 3 last nucleotides downstream of the cleavage site).

**Supplementary Figure 6.**
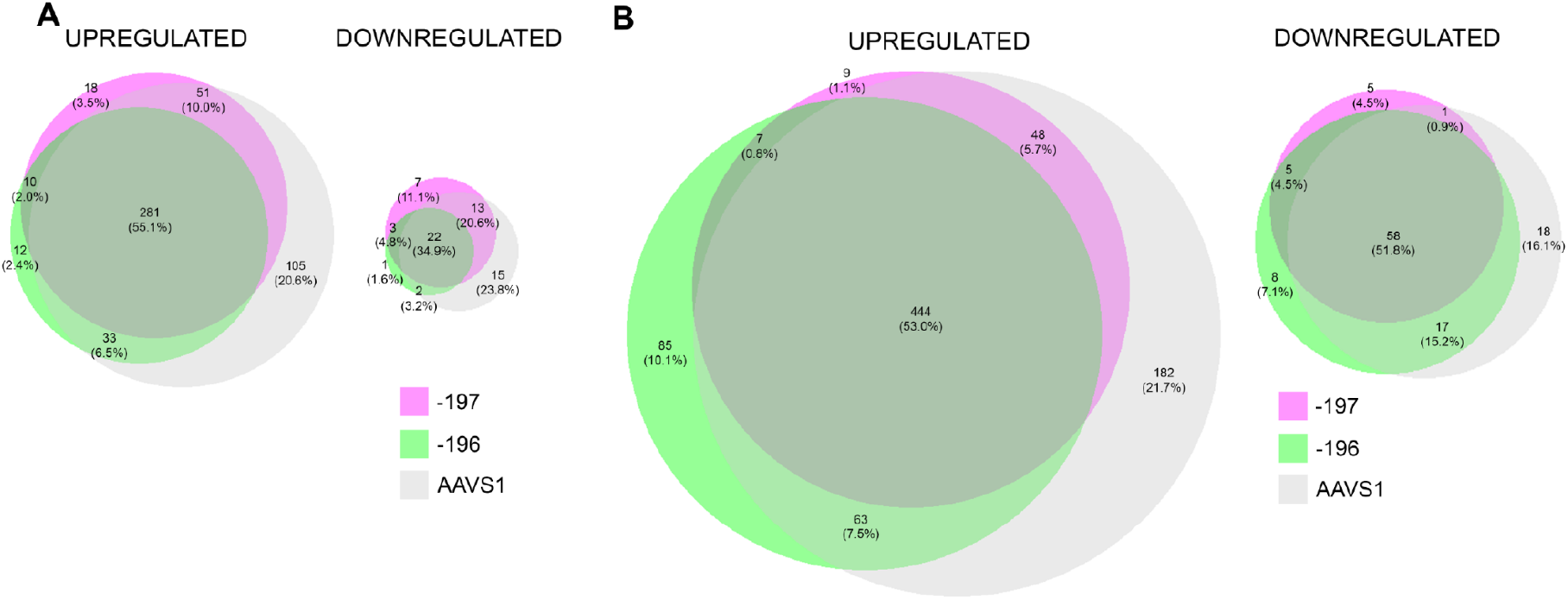
DEGs in primary SCD and HD HSPCs treated with control and *HBG*-targeting sgRNAs. (**A**) Venn diagrams showing the differentially expressed genes (DEGs) upregulated and downregulated in HD samples upon treatment with the sgRNA −197 (pink), −196 (green) and AAVS1 (grey). (**B**) Venn diagrams showing the DEGs upregulated and downregulated in SCD samples upon treatment with the sgRNA −197 (pink), −196 (green) and AAVS1 (grey). In all diagrams, circle size correlates with the number of DEGs. Graphs were generated using BioVenn^63^.

